# Single-cell transcriptomic atlas of human retina from Chinese donors reveals population-specific cellular diversity

**DOI:** 10.1101/2025.03.16.643491

**Authors:** Sen Lin, Yiwen Tao, Luning Yang, Qi Pan, Tengda Cai, Yunyan Ye, Jianhui Liu, Yang Zhou, Quanyong Yi, Zen Haut Lu, Lie Chen, Gareth McKay, Richard Rankin, Yongqing Shao, Weihua Meng

**Affiliations:** Nottingham Ningbo China Beacons of Excellence Research and Innovation Institute, University of Nottingham Ningbo China, Ningbo, China, 315100; Department of Ophthalmology, Lihuili Hospital affiliated with Ningbo University, Ningbo, China, 315040; Department of Cardiology, Lihuili Hospital affiliated with Ningbo University, Ningbo, China, 315040; Ningbo Institute of Innovation for Combined Medicine and Engineering, The Affiliated Li Huili Hospital, Ningbo University, Ningbo, China, 315201; Department of Ophthalmology, The Affiliated Ningbo Eye Hospital of Wenzhou Medical University, Ningbo, China, 315040; PAPRSB Institute of Health Sciences, Universiti Brunei Darussalam, Bandar Seri, Begawan, Brunei Darussalam, BE1410; Centre for Public Health, Institute of Clinical Science, Queen’s University Belfast, Block B, Royal Hospital, Grosvenor Road, Belfast, UK, BT12 6BA; School of Mathematical Sciences, University of Nottingham Ningbo China, Ningbo, China, 315100; Division of Population Health and Genomics, School of Medicine, University of Dundee, Dundee, UK, DD2 4BF; Center for Public Health, Faculty of Medicine, Health and Life Sciences, School of Medicine, Dentistry and Biomedical Sciences, Queen’s University Belfast, Belfast, UK, BT12 6BA

**Author notes:** Corresponding Author: Dr Weihua Meng, Address: Nottingham Ningbo China Beacons of Excellence Research and Innovation Institute, University of Nottingham Ningbo China, Ningbo, China, 315100. Dr Yongqing Shao, Address: Department of Ophthalmology, The Affiliated Ningbo Eye Hospital of Wenzhou Medical University, Ningbo, China, 315040. Sen Lin and Yiwen Tao contributes equally to this paper.

**Keywords:** Single-cell RNA sequencing, Chinese human retina, Cell type and subtype, Cellular functions, Cellular interactions, Cell atlas

## Abstract

The human retina exhibits complex cellular heterogeneity which is critical for visual function, yet comprehensive ethnic-specific references are scarce in ophthalmic transcriptomics. The lack of single-cell RNA sequencing (scRNA-seq) data from Asian populations particularly Chinese donors imposes significant limitations in understanding population-specific retinal biology. We constructed the first comprehensive single-cell transcriptomic atlas of the human retina from Chinese donors, generated through high-throughput scRNA-seq of ∼290,000 viable cells obtained from 18 fresh retinal specimens (living donor and post-mortem specimens). Our multi-level analysis identified 10 distinct retinal cell types, encompassing all major neuronal lineages, Müller glia, astrocytes, microglia. Detailed subcluster analyses further revealed extensive heterogeneity, identifying distinct subtypes within several cell populations such as 7 amacrine cell subtypes and 14 bipolar cell subtypes. Concurrently, through systematic analysis of delineated subtype-specific molecular programs, we mapped their associated biological signaling pathways, functions, and mechanistic processes. This analysis explained the critical involvement of these subpopulations in core biological processes including synaptic organization, neurotransmission, and phototransduction cascades, potentially governing retinal homeostatic regulation and disease mechanisms. Single-cell transcriptomic atlas of the human retina from Chinese donors describes a comprehensive cellular landscape, encompassing major cell types and subtypes including neuronal, glial, and immune populations. This ethnic-specific atlas provided an important reference for understanding retinal development, cellular interaction and disease pathogenesis in Chinese populations, addressing a longstanding gap in ophthalmic transcriptomic resources.

## Introduction

The human retina is a highly specialized neural tissue responsible for detecting light and transmitting visual signals to the brain [1]. This process is mediated by distinct retinal cell types, including rod and cone photoreceptors, bipolar cells (BCs), amacrine cells (ACs), horizontal cells (HCs), and retinal ganglion cells (RGCs), which together form intricate signaling networks [2]. Supporting cells such as Müller glia cells (MGCs), astrocytes, and microglia provide essential metabolic and structural support [3]. Given the complexity of the retina’s cellular composition and function, understanding its molecular landscape at single-cell resolution is essential for elucidating normal visual processing and the pathogenesis of retinal diseases, which are a leading cause of irreversible blindness worldwide [4].

Traditional transcriptomic studies, such as microarrays and bulk RNA sequencing, have provided valuable insights into retinal gene expression. However, these methods analyze pooled RNA from heterogeneous cell populations, masking differences between individual cell types [5]. While previous bulk transcriptomic studies have advanced our understanding of overall retinal gene expression, they lacked the resolution needed to distinguish the distinct molecular signatures of each neuronal and supporting cell subtype [6]. The emergence of single-cell RNA sequencing (scRNA-seq) has transformed retinal research by enabling the identification of individual cell types and their transcriptional profiles at high resolution [7]. This technique has been extensively applied to study the human retina, revealing insights into cellular diversity, gene regulatory networks, and disease-related transcriptional changes.

Several previous studies utilizing scRNA-seq have established detailed atlases of retinal cell types in populations from North America and Europe. A single-cell transcriptomic study identified 18 distinct retinal cell clusters in human donors and implicated *MALAT1* in rod photoreceptor degeneration [8]. Recent scRNA-seq analysis of the human retina reveals that during the transition from normal to advanced age-related macular degeneration, compositional changes are more pronounced in macular rods, microglia, endothelial cells, Müller glia, and astrocytes [9]. Furthermore, scRNA-seq analysis of the mouse retina reveals a distinct subpopulation of Müller glia cells that may play a crucial role in the retinal recovery process following injury [10]. Subsequent scRNA-seq research indicated that Spp1 facilitates microglia-endothelial cell communication and enhances retinal neovascularization, suggesting its potential as a therapeutic target [11]. Despite these progress in retinal research, studies utilizing human samples from Asian populations remain limited, with the majority of retinal atlases derived from America and Europe populations and some mammal models.

Ethic background, environmental factors, and lifestyle habits vary among populations and may influence retinal composition and disease susceptibility [12–15]. South Asians are more susceptible to retinal detachment at a younger age than European Caucasians [14]. Studies have shown that Chinese populations have a higher prevalence of myopia and related retinal diseases, with genetic links between high myopia and retinal disease risk [16]. Additionally, lifestyle factors such as diet, obesity, and physical inactivity have been shown to contribute to the increasing prevalence of retinal diseases in Chinese populations [17]. There is currently a lack of comprehensive single-cell transcriptomic studies that specifically characterize the retinal cell types and subtypes in Chinese individuals.

The present study aims to establish a comprehensive single-cell transcriptomic atlas of the human retina in Chinese population. By analyzing 18 retinal samples comprising 290,401 cells, we seek to identify the cellular diversity, molecular signatures, and subtype-specific biological functions within the Chinese retina, which provides a potential resource for understanding population-specific variations in retinal cell biology.

## Materials and Methods

### Human retinal tissue collection and processing

Collection of human retinal tissues was approved by the ethics committees of the University of Nottingham Ningbo China, The Ningbo Eye Hospital and The Ningbo Medical Centre Lihuili Hospital. Written informed consent was obtained from all living donors. From June 2023 to July 2024, a total of 18 human retinal tissue samples were collected, including two from the Lihuili Hospital and 16 from the Ningbo Eye Hospital. Retinal tissues from six pairs of eyes and one single eye were collected from deceased donors, while the remaining five single-eye samples originated from surgical enucleation procedures. Detailed donor information for all samples is provided in Table 1. Fresh retinal samples in this study refer to those obtained from living donors within 10 minutes after surgical enucleation, as well as those collected from post-mortem donors within 12 hours under temperature-controlled conditions.

**Table 1:** Donor sample information. DM: Samples with diabetes. DR: Samples with diabetic retinopathy. non: Samples without diabetes. SCT-H-PON: Severe craniocerebral trauma with hemorrhage in the periphery of the optic nerve.

| Donor | Eye | Sex | Age | Diabetic Status | Eye Removal/Death Reason |
| --- | --- | --- | --- | --- | --- |
| 01 | Left | M | 55 | Non | Eyeball atrophy |
| 02 | Right | M | 71 | Non | Corneal ulcer |
| 03 | Left | F | 50 | Non | Eyeball atrophy |
| 04 | Right | F | 68 | Non | Corneal perforation |
| 05 | Left | M | 34 | Non | Cerebral hemorrhage |
| 06 | Left | F | 92 | Non | Natural death |
| 07 | Right | F | 92 | Non | Natural death |
| 08 | Left | M | 76 | DM | Corneal staphyloma |
| 09 | Left | F | 55 | DM | SCT-H-PON |
| 10 | Right | F | 55 | DM | SCT-H-PON |
| 11 | Left | M | 55 | DR | Brain stem hemorrhage |
| 12 | Right | M | 55 | DR | Brain stem hemorrhage |
| 13 | Left | M | 35 | DR | Traumatic shock |
| 14 | Right | M | 35 | DR | Traumatic shock |
| 15 | Left | M | 84 | DR | Prostate malignancy |
| 16 | Right | M | 84 | DR | Prostate malignancy |
| 17 | Left | M | 69 | DR | Lung cancer |
| 18 | Right | M | 69 | DR | Lung cancer |

### Single-cell RNA sequencing

Collected human retinal tissue were prepared for scRNA-seq using the Worthington Papain Dissociation Kit (catalog no. LK003150-1bx, Worthington Biochemical Corporation). Tissue samples were stored on ice in a petri plate and chopped into 2–4 mm pieces. The fragments were then incubated in RPMI medium containing the designated enzyme mix (Miltenyi Biotec) at 37°C for 15 minutes. The tissue-enzyme mixture was transferred into a gentleMACS C tube containing additional RPMI/enzyme mix. Following dissociation, the cell mixture was filtered using a 70 µm strainer and centrifuged at 300 × g at 4°C for 10 minutes. The supernatant was discarded, and the pellet was resuspended in 10 volumes of 1× Red Blood Cell Lysis Solution (catalog no. 130-094-183, Miltenyi Biotec) at 4°C for 10 minutes to remove erythrocytes. The suspension was centrifuged again, and the pellet was resuspended in 1x PBS containing 0.04% BSA. The final cell concentration was adjusted to 700-1200 cells/μL, ensuring a viability rate >85%, and assessed using CountStar. Libraries were prepared from 10,000 viable cells per sample using the Chromium Single Cell 3’ Library & Gel Bead Kit (10X Genomics, PN1000268) and subsequently sequenced on the Illumina platform. The quality control step filtered out reads with more than three ‘N’ bases, reads where over 20% of bases had a quality score below 5, and reads containing adapter sequences.

### Generation of single-cell RNA sequencing data

Raw reads were demultiplexed and aligned to the reference genome (GRCh38-2020-A) using the 10X Genomics Cell Ranger with default parameters (https://www.10xgenomics.com/support/software/cell-ranger). Cell Ranger handles FASTQ files by performing tasks including alignment, filtering, barcode counting, and UMI quantification. It utilizes Chromium cellular barcodes to build feature-barcode matrices, which are subsequently used for dimensionality reduction, clustering, and gene expression analysis based on Cell Ranger’s algorithms.

### Quality control and doublets removal

After the raw data were processed by using 10X Genomics Cell Ranger pipeline, the standard 10X output files that are barcodes, features and matrix were obtained. We used some quality control methods to filter out uninformative, outlier and error data. Based on the calculation results of gene expressions and mitochondrial read fraction, we defined the criteria of high-quality cell data: the number of genes expressed in each cell was greater than 50, the total counts per cell were greater than 400, and the percentage of counts in mitochondrial genes was less than 10%. Low-quality cell data were filtered out. To further improve the quality of data, cell doublets in data were identified and removed using the doublet detection method Scrublet [18] in Python which identified cell doublets through a nearest-neighbor classifier of simulated doublets and observed transcriptomes.

### Batch effect and data integration

High-quality data for downstream analysis was selected in each sample after quality control. We have 18 samples from different donors. Batch effect can be one of the non-negligible factors lead to misclassifications or distortions in downstream analysis steps. We applied Batch Balanced K Nearest Neighbours (BBKNN) to remove batch effect. BBKNN could identify each cell’s k nearest neighbors from each batch and combine its nearest neighbors into the final neighbors for the cell [19]. This method BBKNN clusters similar cells from different samples together, thereby effectively eliminating the impact of different samples on subsequent analysis.

### Data normalization and scaling

We used the Python toolkit Scanpy (v1.10.1) [20] to complete the following data preprocessing steps. In scRNA-seq data, the total counts of transcripts vary across different cells due to the influence of some technical bias such as sequencing depth, cell capture efficiency, and library construction. The data was normalized and scaled with log plus one (log1p) transformation. The normalization process helps eliminate technical bias between cells so that different cells could be comparable in single cell gene expression levels. The difference of gene expression counts often results in a strongly skewed distribution in raw data, which affects subsequent preprocessing steps such as dimensionality reduction, cell clustering, and differential expression analysis. We performed log transformation to make the data distribution smoother and more in line with the assumptions of downstream methods (e.g. t-SNE, UMAP) on the data distribution. After data normalization and scaling, the scRNA-seq raw data was adjusted to the same numerical scale, which not only mitigates the effects of technical bias, but also ensures the downstream data analysis more directly reflect the real differences in biology.

### Data dimensionality reduction and cell clustering

Single-cell transcriptomes technique usually measure a very large number of genes, which could contain a lot of genes whose expression is useless, stable or noisy. These genes may contribute little to distinguishing cell types while adding dimensionality and noise to the data. After data preprocessing, we selected the top 5,000 most variable genes for the next analysis steps. The expression levels of these most variable genes always fluctuate greatly across different cells, which can precisely reflect changes and differences in cell biological identity and state. While ensuring the integrity of biological information, the dimensionality of data is greatly reduced, and the interference of noise is also reduced.

We ran running principal component analysis (PCA) to reduce the dimensionality of data. Through the linear transformation, high-dimensional data can be projected into low-dimensional space. These principal components are able to capture most of the biological variation information in the raw data, thus retaining the most useful biological signals and ignoring noisy and redundant information.

We applied the Leiden graph-clustering method to partition cells into distinct, biologically meaningful groups [21]. This graph-based approach is particularly effective for scRNA-seq data. The resolution parameter of the Leiden algorithm was varied from 0.02 to 2.00 to identify and analyze different cell populations. For each iteration, a K-Nearest Neighbors (KNN) graph was generated at each resolution parameter value. We selected a reasonable and clear KNN graph and used UMAP to visualize the cell clusters in two-dimensional space, facilitating the interpretation of the biological significance of each cluster.

### Cell type annotation

Based on the UMAP visualization, we manually annotated the cell clusters to delineate distinct subpopulations by using known marker genes. The UMAP provided a robust, two-dimensional representation of the cells, which could visually identify discrete clusters that correspond to unique cell types or cellular states. By integrating the prior biological knowledge with the expression patterns of canonical marker genes, we were able to assign cell type to each cell cluster.

### Differential expression analysis

After cell type annotation, we used the FindAllMarkers function in R (v4.4.1) Seurat package (v5.2.0) to perform statistical tests to find genes that are differentially expressed across various human retinal cell types and subtypes [22]. Both upregulated and downregulated genes were considered, with the requirement that these genes must be expressed in at least 20% of the cells in a cell cluster. The statistical test method Wilcoxon Rank Sum test was applied for scRNA-seq data. The average log2 fold change represents the average difference in gene expression between cell clusters, transformed on a logarithmic scale, which determines whether the gene is upregulated or downregulated, with values greater than 0 being upregulated and values less than 0 being downregulated. For further research, we labelled the genes with an absolute value of average log2 fold change greater than 1 as highly fold changes and those with an absolute value less than 1 as lowly fold changes in the results. Given the biological context and relevance, even smaller differences in gene expression may be biologically significant, and it is important to consider both smaller and larger differences to draw meaningful conclusions. We performed differential expression analysis on each human retinal cell type and subtype, which reveals cellular heterogeneity and helps to identify key regulatory genes and pathways, crucial for understanding human retinal complexity.

### Enrichment analysis

Gene Ontology (GO) enrichment analysis identified biological processes in a set of genes, resulting from differential expression analysis [23]. GO enrichment analysis allowed us to recognize a list of genes with specific biological functions, processes, or cellular components, providing insights into the biological mechanisms underlying a particular cell condition or type. The clusterProfiler package (v4.12.6) in R was used to perform GO enrichment analysis. It tests whether certain BP-related GO terms are statistically enriched in the input gene set compared to the genome-wide background by using the human database containing gene information for Homo sapiens. The adjusted p-value was used as a significance threshold for GO term enrichment, which only GO terms having an adjusted p-value less than 0.05 are considered statistically significant.

Kyoto Encyclopedia of Genes and Genomes (KEGG) pathway enrichment analysis was used to identify biological pathways in a set of genes, derived from differential expression analysis [24]. By linking genes to specific KEGG pathways, KEGG pathway enrichment analysis helps to understand key biological mechanisms associated with developmental processes and biological phenomena. The clusterProfiler package in R is used to perform KEGG pathway enrichment analysis. The Benjamini-Hochberg (BH) method was applied to the adjust p-value for multiple tests, reducing the false discovery rate (FDR) and ensuring the results are statistically valid. The q-value is a significance threshold for KEGG pathway enrichment analysis, where KEGG pathways with value less than 0.05 are considered statistically significant.

### Cell-cell communication network analysis

Cell-cell communication network analysis is a crucial aspect of understanding cellular interactions in the levels of single-cell transcriptomics [25]. The CellChat package (v2.1.2) in R was a powerful tool used for inference of cell-cell interactions, key signaling pathways identification, and cell-cell communication networks visualization using the single-cell transcriptomic data from individual cells. It identifies signaling pathways and interactions between cells by integrating receptor-ligand interactions and signaling pathways from the database CellChatDB. The database CellChatDB contains information on cell-to-cell signaling pathways in a variety of human organisms. The method “triMean” was used to compute the average gene expression per cell subtype and the minimum number of cells required in each cell cluster is 10, which could produce fewer but stronger interactions. By using the tool CellChat, we analyzed the gene expressions of receptors and ligands in different cell types, identify which cell types are likely involved in specific signaling interactions and quantify the strength of each signaling pathway and receptor-ligand pair across different cell types.

## Results

### Construction of human retinal scRNA-Seq cell atlas

We constructed and analyzed comprehensive human retinal single-cell transcription cell atlas from 18 Chinese human retinal samples that were collected from living donors and postmortem specimens. After data was preprocessed following the procedures, we visualized cell clusters across all samples in the UMAP (Figure 1). Through comprehensive single-cell transcriptomic analysis, we meticulously characterized the cellular heterogeneity of the retina, identifying 10 distinct cell populations: cone photoreceptors marked by *ARR3*, *GNAT2*, and *GNGT2*; rod photoreceptors marked by *RHO*, *PDE6G*, *GNGT1*, *NRL*; BCs marked by *GRM6* and *GRIK1*; ACs marked by *TFAP2B*, *GAD1* and *SLC32A1*; astrocytes marked by *GFAP*; MGCs marked by *RLBP1*, *RGR*, and *DKK3*; RGCs marked by *RBPMS* and *SLC17A6*; HCs marked by *LHX1* and *TNR*; microglia marked by *CD74*, and *TYROBP*; and T cells marked by *CD69*, *CD52*, and *CD3D* (Figure 2A) [26].

**Figure 1:**
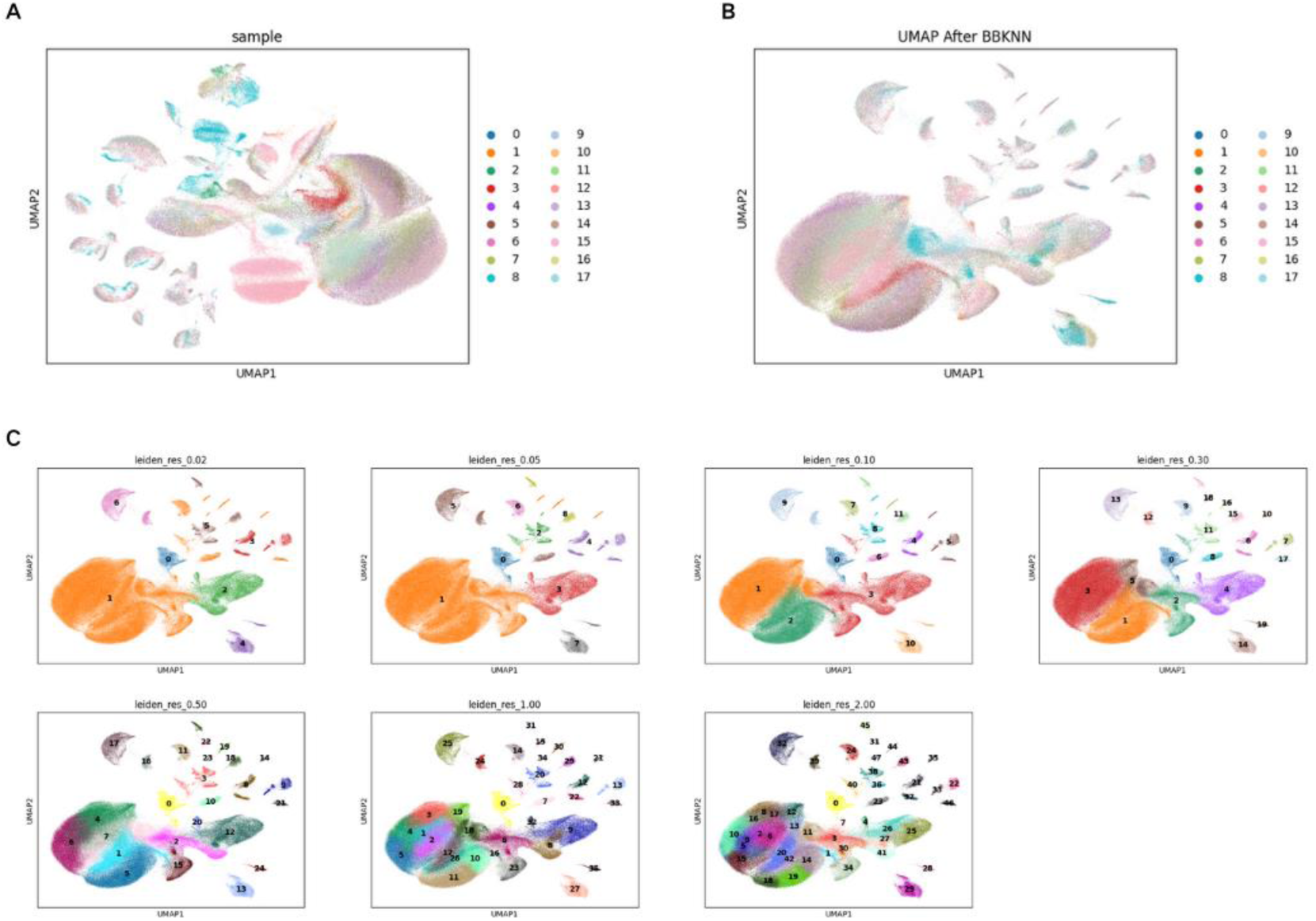
batch effect removal and cell clustering. (A) UMAP plotted of scRNA-seq data integrated from 18 human retina samples. Each dot represented a cell. The distribution of cells was not evenly but very influenced by batch effects caused by different samples. (B) UMAP of scRNA-seq data integrated from 18 human retina samples after batch effect removal by using BBKNN. After BBKNN, the distribution was not influenced by sample. (C) UMAP plotted illustrated the cell clusters by using Leiden algorithm resolutions at various resolutions, from 0.02 to 2.00. We chose the clustering resolution of 1.00 as the best value, because even for some cell types with a small percentage of cells, they can be clearly and accurately annotated.

**Figure 2:**
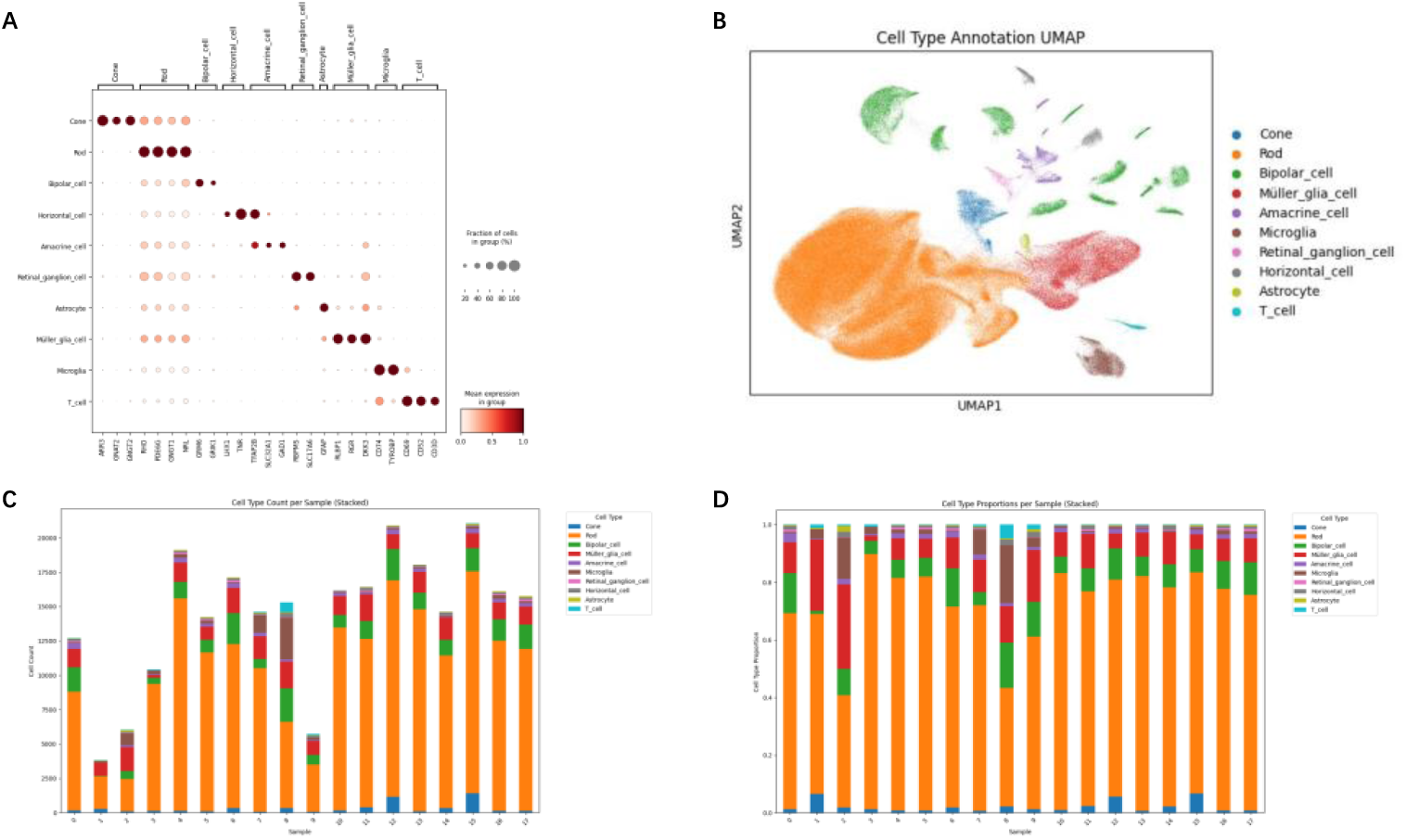
Comprehensive analysis of Chinese human retinal cell types and clusters. (A) Dot plot demonstrated the identification of each cluster using known markers of the major cell types, and the selection of genes that differentiated each cluster in retinal samples. Each row corresponds to a specific major cell type, and the columns correspond to the gene expressions of a set of key marker genes. At the top of the plot detailed key marker genes for each cell type. (B) UMAP plotted that each cell cluster was annotated by major human retinal cell types. (C) Stacked bar chart plotted the number of each cell type in each human retinal sample. (D) Stacked bar chart plotted the proportion of each cell type in each human retinal sample.

After cell cluster annotation, we gained a comprehensive understanding of the composition of cell types in the human retina. The total number of preprocessed cells for downstream analysis was 258,164 from 18 Chinese donor samples (Figure 2B). The predominant cell population was comprised of rod photoreceptors, accounting for 73.3% of the total cells. MGCs, provides structural support to maintain retinal stability and integrity, represented 9.4% of the population. BCs, which play a key role in transmitting visual information from photoreceptors to retinal ganglion cells, made up 8.9%. A smaller proportion of the retina was composed of microglia (2.8%), cone photoreceptors (2.1%), and ACs (1.5%), which are involved in modulating retinal signaling. HCs, important for visual information processing, represented 0.9%, while T cells (0.4%), RGCs (0.4%), and astrocytes (0.3%) were less abundant. The human retinal scRNA-Seq cell atlas reveals the cellular composition of the human retina through single-cell transcriptomics (Figure 2B, 2C and 2D).

### Identification and Analysis of Human Retinal Cell Subtype

A diverse array of cell types and their respective proportions was presented in the human retinal scRNA-Seq cell atlas we constructed (Figure 1). To further investigate the subtypes within each cell population, we perform the same cell clustering and cell type annotation procedures on each cell type. Then we identified subtypes of each human retinal cell type by analyzing the expression profiles of specific marker genes within each major cell subtype. These distinct cell subtypes or subgroups that may represent functionally specialized or developmentally distinct populations within the broader retinal cell types based on bioinformatic analysis.

### Amacrine Cells

Amacrine cells (ACs) are a diverse cell cluster of interneurons in the human retina, playing a crucial role in modulating retinal signal processing and contributing to visual information integration [27]. In general, ACs could be classified to three major subtypes GABAergic, glycinergic, or neither, which can be determined by the inhibitory neurotransmitter they express (GABA, glycine, or neither) [28–29]. We identified 7 subtypes of amacrine cell subtypes that were Sodium-Dependent Glutamate Transporter-

Expressing Amacrine Cells (SEG-ACs), Substance P-Expressing Amacrine Cells (SubstanceP-ACs), Starburst Amacrine Cells (Starburst ACs), CART Peptide-Expressing Amacrine Cells (CARTPT-ACs), AII amacrine cells (All ACs), Neuropeptide Y-Expressing Amacrine Cells (NPY-ACs), Vesicular Glutamate Transporter 3-Expressing Amacrine Cells (VG3 ACs) in our dataset (Figure 3B). The known gene markers for identification were: *PVALB*, *SATB2*, and *EBF3* for SEG-ACs; *TACR1* and *TACR3* for SubstanceP-ACs; *MEGF10* and *TENM3* for Starburst ACs; *CARTPT* for CARTPT-ACs; *GJD2* and *CALB2* for All ACs; *NPY* for NPY-ACs; *SLC17A8* for VG3 ACs. GABAergic ACs contains SubstanceP-ACs, Starburst ACs, CARTPT-ACs, and NPY-ACs, while glycinergic ACs includes SEG-ACs, All ACs, and VG3 ACs (Figure 3A) [30].

**Figure 3:**
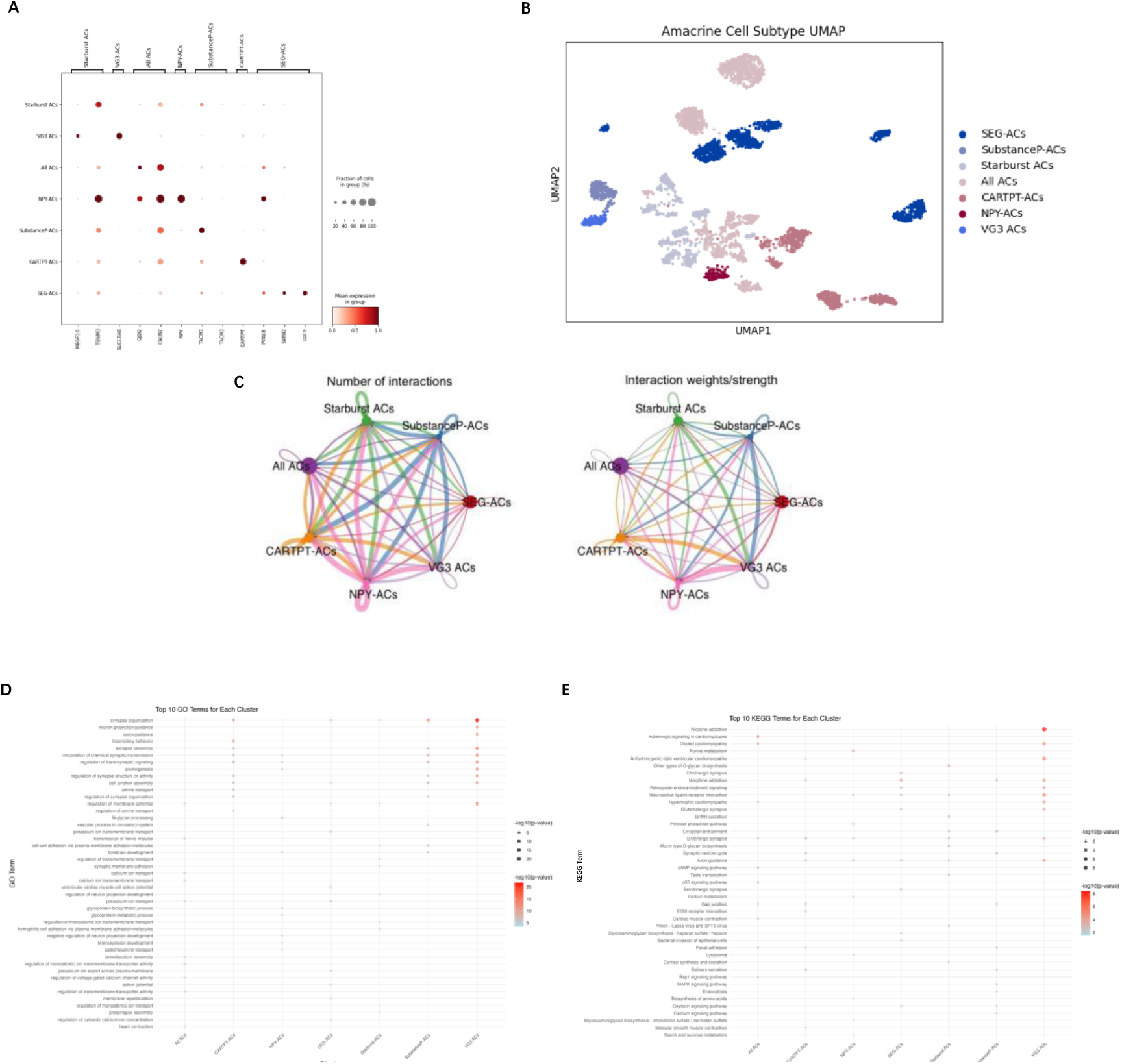
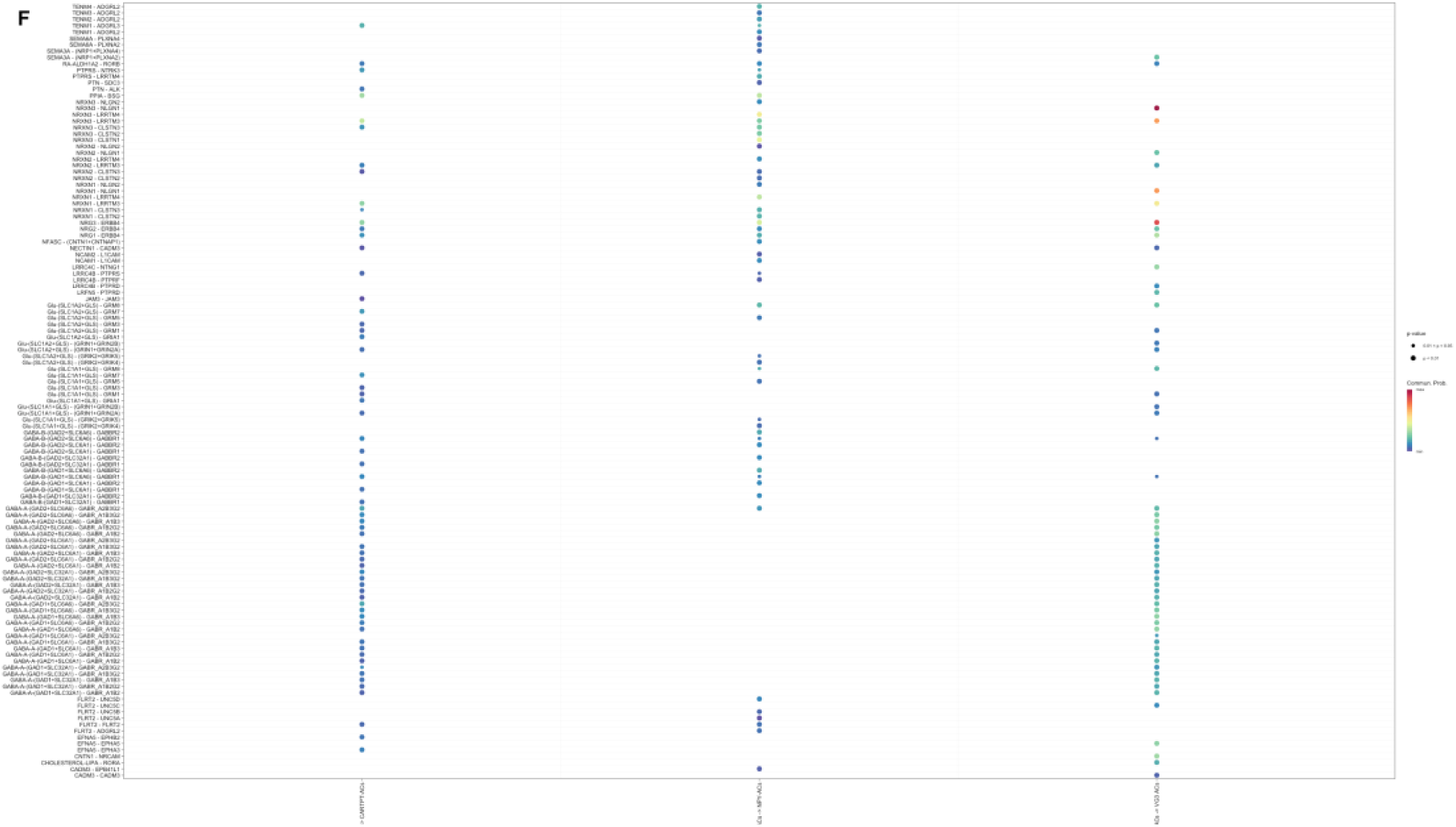
Comprehensive analysis of Chinese human retinal ACs type and subtypes. (A) Dot plot demonstrated the identification of each cluster using known markers of subtypes of ACs. (B) UMAP plotted that each cell cluster was annotated by ACs subtypes. (C) Circle plot visualized the aggregated cell-cell communication network that presented the number of interactions (the left) and the total interaction strength/weights (the right) between any two cell subtypes. (D) Dot plot presented the top 10 enriched GO terms for each cell subtype. The row represented each GO term that these subtypes were involved in. Dots represented that cell subtypes were associated with the GO term, the redder the color, the deeper the correlation. (E) Dot plot presented the top 10 enriched KEGG terms for each cell subtype. The row represented each KEGG term that these subtypes were involved in. Dots represented that cell subtypes were associated with the KEGG term, the redder the color, the deeper the correlation (F) Bubble plot showed all the significant interactions mediated by ligand-receptor pairs and signaling pathways from some cell subtypes to other cell subtypes. The row represented the significant interactions (ligand-receptor pairs) and signaling pathways between two cell subtypes. Dots represented the related interactions and signaling pathways, the redder the color, the deeper the correlation.

In bioinformatic analysis, the functions of SEG-ACs are involved in the regulation of glutamate levels within the retina, help maintain the balance of excitatory neurotransmission and modulates visual signal processing in the retina. SubstanceP-ACs release substance P and participate in neurotransmission and modulate synaptic activity. Starburst ACs release acetylcholine and are involved in processing visual information related to motion detection and directionality [31]. CARTPT-ACs express the cocaine- and amphetamine-regulated transcript (CART) peptide, which are involved in modulating neurotransmission and regulating circadian rhythms and energy homeostasis within the human retina. All ACs are used to assist in the interpretation of photoreceptor signals and transmit light signal from rod cells to retinal ganglion cells. NPY-ACs release neuropeptide Y involved in various physiological processes and significate for synaptic activity, retinal transmission, retina pathophysiology and stress reaction. VG3 ACs express vesicular glutamate transporter 3 (VGLUT3), which are involved in excitatory neurotransmission within the retina, contributing to the processing of visual information and the regulation of circadian rhythms.

GO enrichment analysis listed the GO terms such as “synapse organization,” “axon guidance,” and “modulation of chemical synaptic transmission,” which are significant to the neural and synaptic activities of these cell subtypes of ACs (Figure 3D). These processes are important for understanding how ACs contribute to retinal signaling. KEGG enrichment analysis provides a deeper understanding of the metabolic, signaling, and physiological processes in ACs (Figure 3E). The included terms such as “Nicotine addiction,” “Adrenergic signaling in cardiomyocytes,” and “Axon guidance,” which displayed these cells were involved in synaptic transmission, diseases and signal processing in the retina. The results of CellChat showed that the cell subtypes of ACs associated with the pathways neurexins (NRXN1, NRXN2, NRXN3), GABAergic (GABA-B and GABA-A), and glutamatergic pathways (GLU), which played an important role in the synaptic organization, cell adhesion, synaptic signaling and neuronal regulation, and neurodevelopmental processes (Figure 3C and 3F).

### Bipolar Cells

Bipolar cells (BCs) locate in the inner nuclear layer of the retina. Served as intermediaries, BCs transmit visual information and signal from photoreceptors (rod photoreceptors and cone photoreceptors) to RGCs. Structurally, BCs are characterized by a bipolar shape, with receiving synaptic input from photoreceptors and axons synapsing with RGCs. Functionally, they play a crucial role in processing visual information and signals, contributing to the retina’s ability to detect light intensity and contrast, especially in dark and light environments [32].

We observed 14 subtypes of BCs in our data using known gene markers (Figure 4B). They are listed separately: *LRPPRC* for diffuse bipolar 1 (DB1); *AFA4* for diffuse bipolar 2 (DB2); *CALB1* for diffuse bipolar 3a (DB3a); *MEIS2* for diffuse bipolar 3b (DB3b); *TTR* for diffuse bipolar 4a (DB4a); *PRSS12* for diffuse bipolar 4b (DB4b); *DOK5* for diffuse bipolar 5 (DB5); *NELL2* for diffuse bipolar 6 (DB6); *FEZF1* for OFFx; *SCG2* for flat midget bipolar (FMB); *ODF2L* for invaginating midget bipolar (IMB); *AGBL1* for giant bipolar (GB); *SORCS3* for blue bipolar (BB); *PRKCA* and *NIF3L1* for Rod bipolar cells (RB) (Figure 4A) [33–34]. These cell subtypes could be classified to a higher level in two ways [35]. One way is rod versus cone BCs: only RB belongs to rod BCs; all the remaining cell subtypes belong to cone BCs. The second way is ON versus OFF BCs: ON BCs include DB4a, DB4b, DB5, DB6, IBM, BB, GB and OFF BCs include DB1, DB2, DB3a, DB3b, FMB, OFFx.

**Figure 4:**
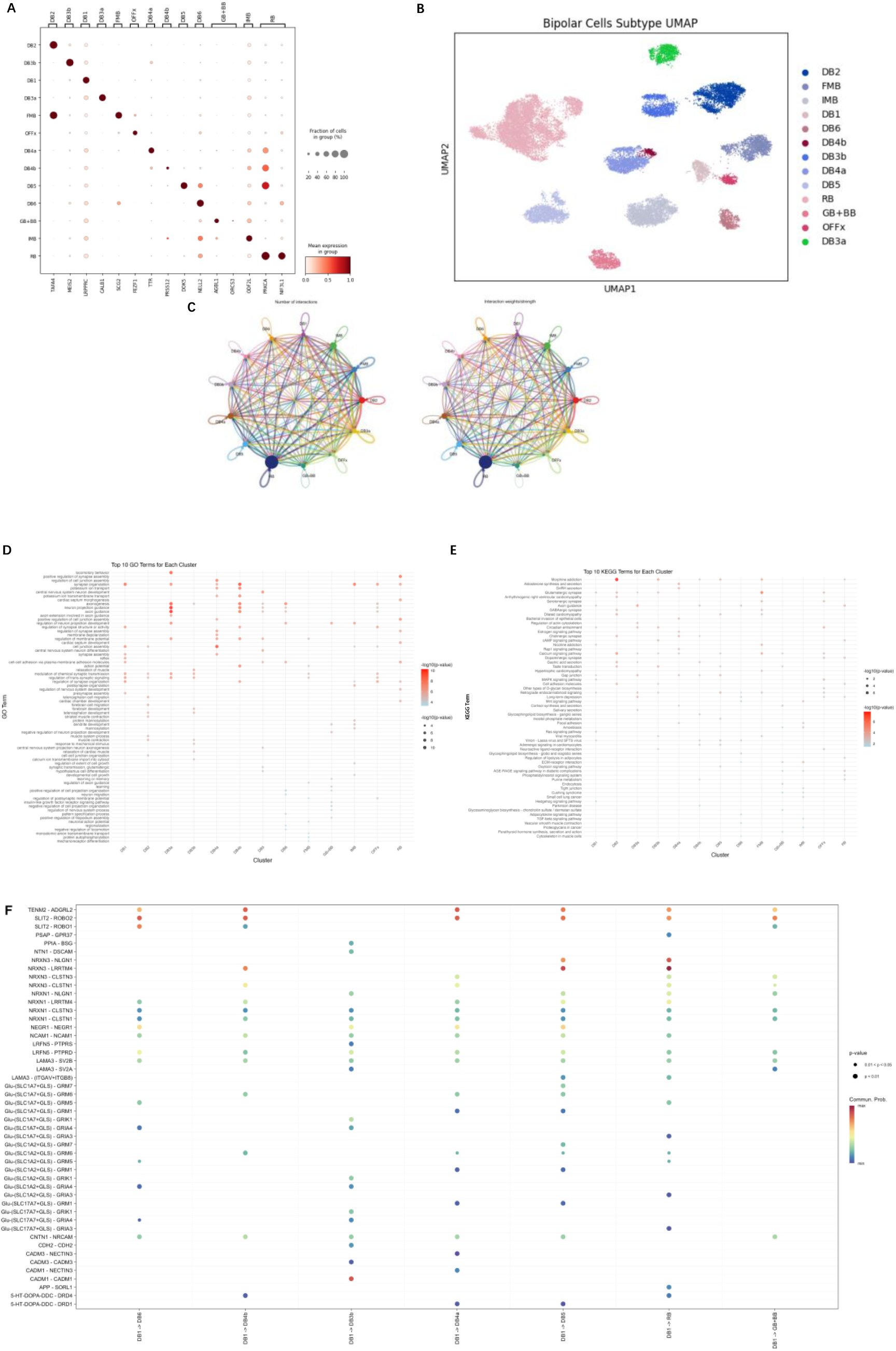
Comprehensive analysis of Chinese human retinal BCs type and subtypes. (A) Dot plot demonstrated the identification of each cluster using known markers of subtypes of BCs. (B) UMAP plotted that each cell cluster was annotated by BCs subtypes. (C) Circle plot visualized the aggregated cell-cell communication network that presented the number of interactions (the left) and the total interaction strength/weights (the right) between any two cell subtypes. (D) Dot plot presented the top 10 enriched GO terms for each cell subtype. The row represented each GO term that these subtypes were involved in. Dots represented that cell subtypes were associated with the GO term, the redder the color, the deeper the correlation. (E) Dot plot presented the top 10 enriched KEGG terms for each cell subtype. The row represented each KEGG term that these subtypes were involved in. Dots represented that cell subtypes were associated with the KEGG term, the redder the color, the deeper the correlation (F) Bubble plot showed all the significant interactions mediated by ligand-receptor pairs and signaling pathways from some cell subtypes to other cell subtypes. The row represented the significant interactions (ligand-receptor pairs) and signaling pathways between two cell subtypes. Dots represented the related interactions and signaling pathways, the redder the color, the deeper the correlation.

Both ON and OFF BCs are two primary neurons in the human retina, which performs a crucial role in processing visual information and signals [36]. The axon terminals of ON BCs are located mainly in the inner layer of the inner plexiform layer of the retina, while the axon terminals of OFF BCs are in the outer layer of the inner plexiform layer. This layered structure allows the retina to efficiently process increases and decreases in light, creating contrast-sensitive visual signals. ON and OFF BCs are connected to photoreceptors (rod and cone photoreceptors) and are regulated by glutamate [37–38]. ON BCs depolarize in response to light increments and hyperpolarize when light diminishes. In contrast, OFF BCs depolarize when light decreases and hyperpolarize when light increases. In darkness, photoreceptors continuously release glutamate to hyperpolarize ON BCs and decrease OFF BCs. With light exposure, reduced glutamate release leads to hyperpolarization of OFF BCs and depolarization of ON BCs.

GO enrichment analysis showed these subtypes of BCs were involved in neuronal development and synaptic regulation such as regulation of synaptic signaling, central nervous system development, ion transport and signaling like calcium ion and negative regulation and homeostasis. (Figure 4D). The KEGG enrichment analysis results suggested that BCs subtypes engage in diverse molecular processes, including neurotransmission, metabolic regulation, synapse activity and disease-associated pathways. (Figure 4E). Communication strength between BCs subtypes was quantified (Figure 4C). These ligand-receptor pairs showed that the subtypes were involved in neuronal connectivity (such as GLU, TENM2 and ADGRL2), neurotransmitter (NRXN3), organogenesis, disease mechanisms and visual signal processing (SLIT2 and ROBO2/ROBO1) (Figure 4F).

The distinct responses of ON and OFF BCs enable the human retina to process visual information such as contrasts effectively, facilitating the detection of light and dark environments within the visual field.

### Retinal Ganglion Cells

Retinal ganglion cells (RGCs) located in the ganglion cell layer of the human retina, near the inner surface of the eye [39]. The function of RGCs is to receive synaptic inputs from BCs and ACs, which process visual signals from photoreceptors (rod and cone photoreceptors). The axons of RGCs converge to form the optic nerve, which transmits visual information to various brain regions such as hypothalamus and midbrain [40]. RGCs play a crucial role in visual processing by integrating and transmitting information from the human retina to the brain.

Three major subtypes of RGCs that are midget RGCs, parasol RGCs and intrinsically photosensitive RGCs were identified in our data (Figure 5B). The known cell markers are: *TBR1* for OFF midget RGCs; *TPBG* for ON midget RGCs; *FABP4* for OFF parasol RGCs; *CHRNA2* for ON parasol RGCs and *OPN4* for Intrinsically photosensitive RGCs (Figure 5A) [33].

**Figure 5:**
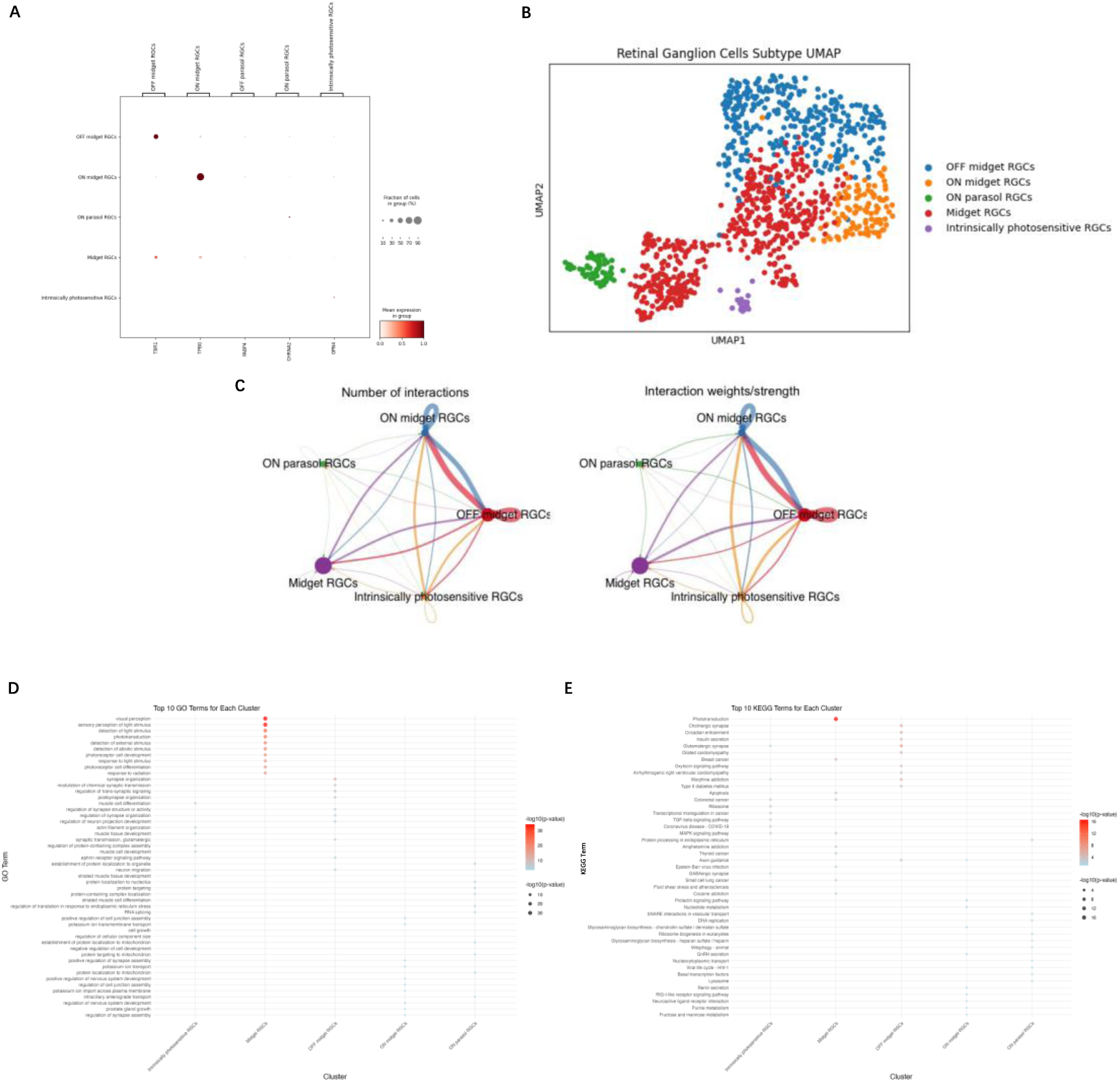
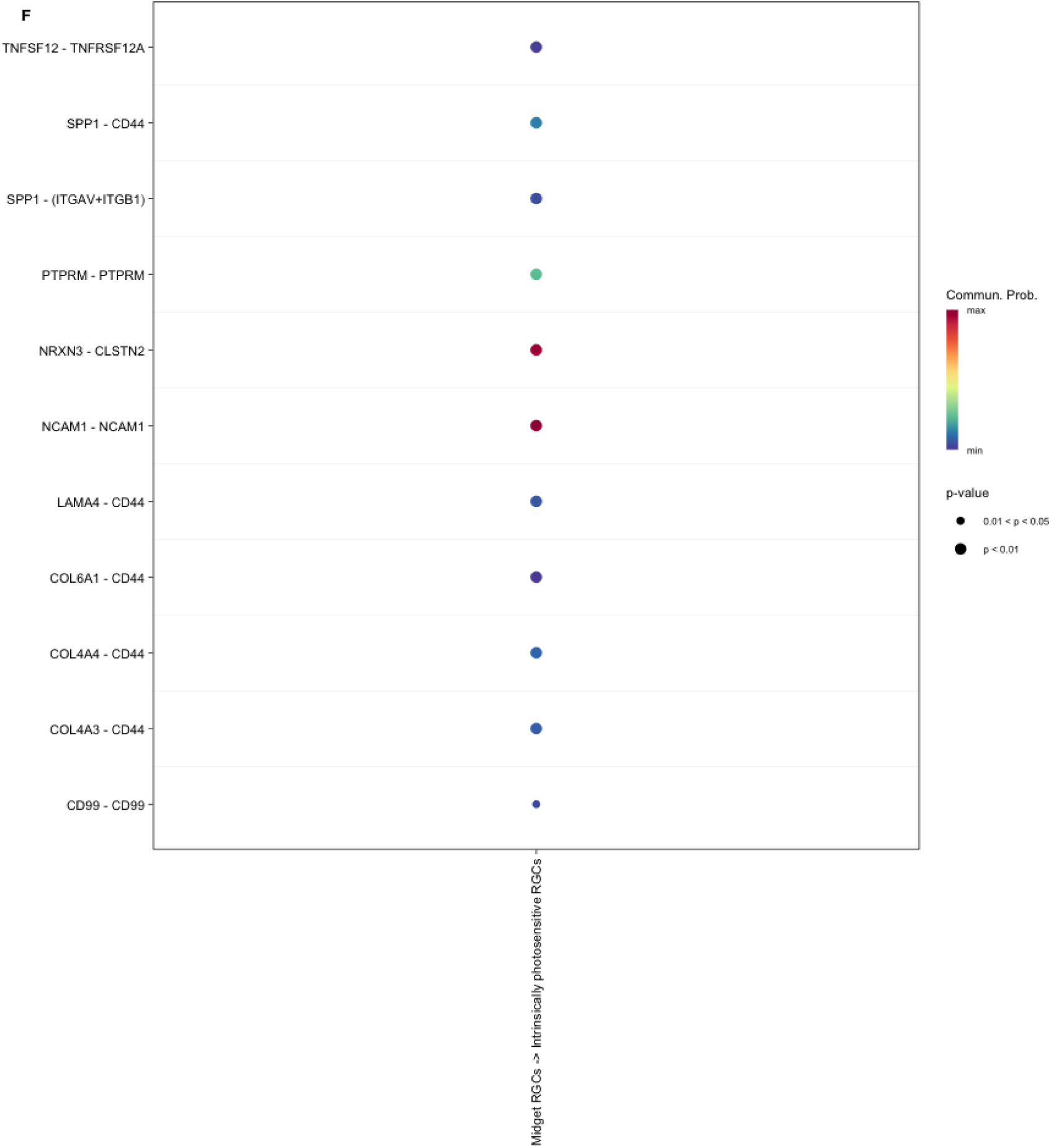
Comprehensive analysis of Chinese human retinal RGCs type and subtypes. (A) Dot plot demonstrated the identification of each cluster using known markers of subtypes of RGCs. (B) UMAP plotted that each cell cluster was annotated by RGCs subtypes. (C) Circle plot visualized the aggregated cell-cell communication network that presented the number of interactions (the left) and the total interaction strength/weights (the right) between any two cell subtypes. (D) Dot plot presented the top 10 enriched GO terms for each cell subtype. The row represented each GO term that these subtypes were involved in. Dots represented that cell subtypes were associated with the GO term, the redder the color, the deeper the correlation. (E) Dot plot presented the top 10 enriched KEGG terms for each cell subtype. The row represented each KEGG term that these subtypes were involved in. Dots represented that cell subtypes were associated with the KEGG term, the redder the color, the deeper the correlation. (F) Bubble plot showed all the significant interactions mediated by ligand-receptor pairs and signaling pathways from some cell subtypes to other cell subtypes. The row represented the significant interactions (ligand-receptor pairs) and signaling pathways between two cell subtypes. Dots represented the related interactions and signaling pathways, the redder the color, the deeper the correlation.

Midget RGCs are predominantly located in the human central retina and responsible for high-acuity vision [41]. They specialize in fine visual detail and spatial resolution, essential for color vision and high-resolution visual processing [42]. ON midget RGCs are activated when light intensity increases, and OFF midget RGCs are activated when light intensity decreases. So, they facilitate contrast detection in both bright and dark conditions. Parasol RGCs are located more peripherally within the human retina. They are the key components of the human retina’s magnocellular pathway, which processes motion and broad visual features. Similarly, ON midget RGCs and OFF midget RGCs work in bright and dark conditions, respectively. intrinsically photosensitive RGCs are found across the human retina. They are primarily involved in regulating non-image-forming visual functions such as circadian rhythms, pupil constriction, light adaptation, and body’s internal clock.

Based on the results of GO and KEGG enrichment analysis, these cell subtypes of RGCs associated with phototransduction, visual perception and sensory perception and detection of light stimulus (Figure 5D and 5E). The result of CellChat showed these subtypes involved in diverse biological processes, ranging from cell adhesion to immune regulation, associated with modulates, including neural development (NCAM1), neuronal migration, axon guidance and immune synapse formation (CD99) (Figure 5C and 5F).

### Horizontal Cells

Horizontal cells (HCs) are inhibitory interneurons located in the retina, receiving input from photoreceptors (rod and cone photoreceptors) and form lateral connections with BCs [42–43]. This lateral interaction could enhance contrast sensitivity and edge detection in visual perception. By modulating the activity of neighboring neurons, HCs facilitate the processing of visual information and signals and improve the clarity and contrast of the visual scene.

We identified 2 subtypes of HCs that are H1 and H2 HCs (Figure 6B). The known cell markers we used are *LHX1* for H1 HCs and *PCDH11X*, *CHN1*, and *ISL1* for H2 HCs [30] (Figure 6A). H1 HCs receives inputs from L cone photoreceptors and M cones photoreceptors and H2 HCs receives strong inputs from S cone photoreceptors and weaker inputs from the other cone photoreceptors types.

**Figure 6:**
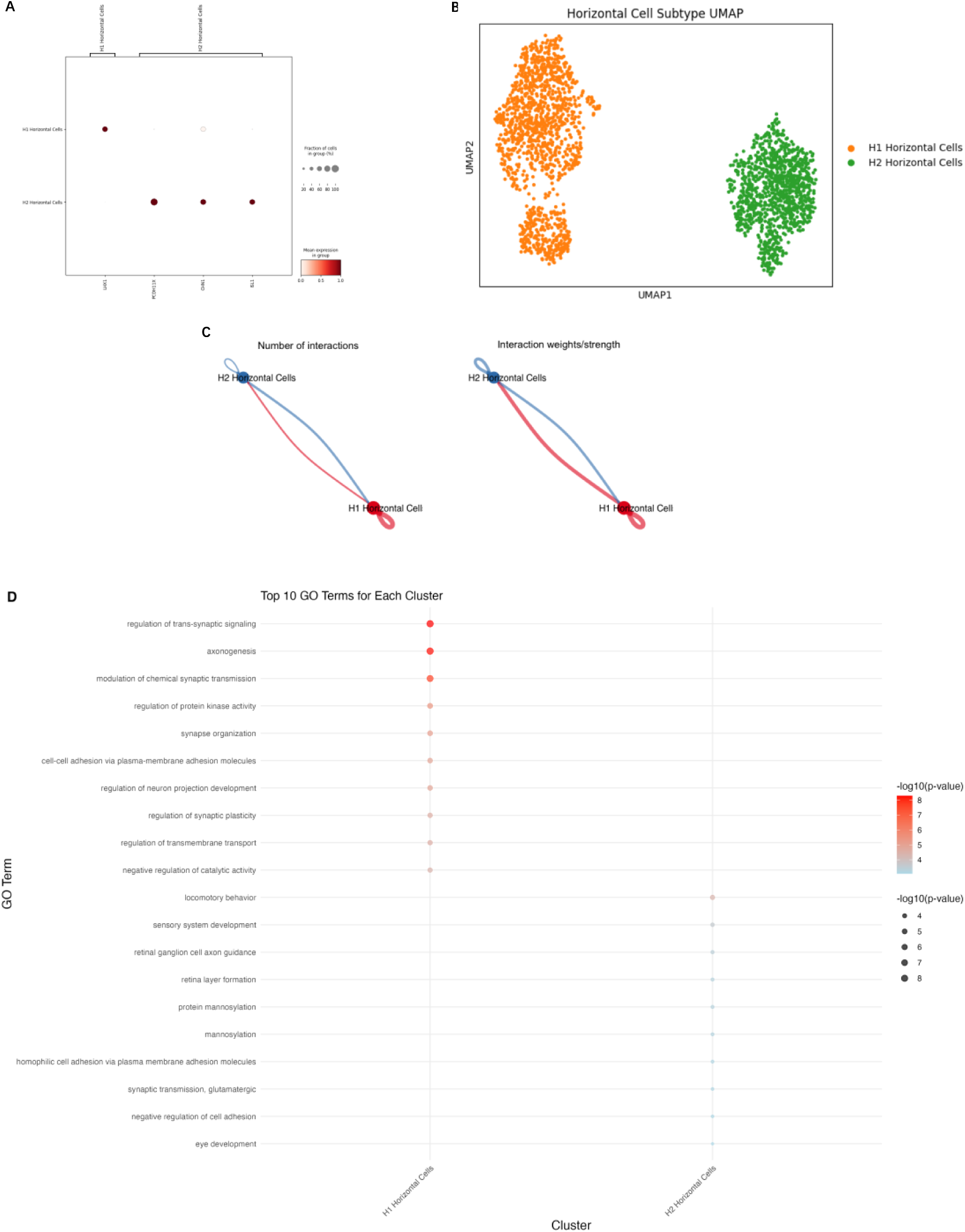
Comprehensive analysis of Chinese human retinal HCs type and subtypes. (A) Dot plot demonstrated the identification of each cluster using known markers of subtypes of HCs. (B) UMAP plotted that each cell cluster was annotated by HCs subtypes. (C) Circle plot visualized the aggregated cell-cell communication network that presented the number of interactions (the left) and the total interaction strength/weights (the right) between any two cell subtypes. (D) Dot plot presented the top 10 enriched GO terms for each cell subtype. The row represented each GO term that these subtypes were involved in. Dots represented that cell subtypes were associated with the GO term, the redder the color, the deeper the correlation.

GO terms indicated that H1 and H2 HCs were associated with regulation of trans-synaptic signaling, axonogenesis, and modulation of chemical synaptic transmission (Figure 6D). As for KEGG enrichment analysis, no significant enriched pathways were detected. The CellChat analysis showed no interactions are detected.

### Cone Photoreceptors

Cone photoreceptors are specialized photoreceptor cells that are most densely concentrated in the central region of human retina, essential for perceiving a wide range of colors and fine detail [44]. The functions of cone photoreceptors are primarily responsible for color vision and function optimally under bright light conditions. Structurally, photopigments in the cone photoreceptors absorb light, produce chemical changes, generate electrical signals, then transmit to the brain via the optic nerve, finally result in visual perception.

There are 3 subtypes of cone photoreceptors, each sensitive to different wavelengths of light [45] (Figure 7B). We used known cell markers to identify these three subtypes: *OPN1SW* for S cone photoreceptors; *OPN1MW* for M cone photoreceptors; and *OPN1LW* for L cone photoreceptors (Figure 7A). S cone photoreceptors are sensitive to short wavelengths, corresponding to blue light. M cone photoreceptors are sensitive to medium wavelengths, corresponding to green light. L cone photoreceptors are sensitive to long wavelengths, corresponding to red light. The combined input from these three subtypes of cone photoreceptors corresponding to three colors blue, green and red enables the human visual system to perceive a broad spectrum of colors through a process known as color vision.

**Figure 7:**
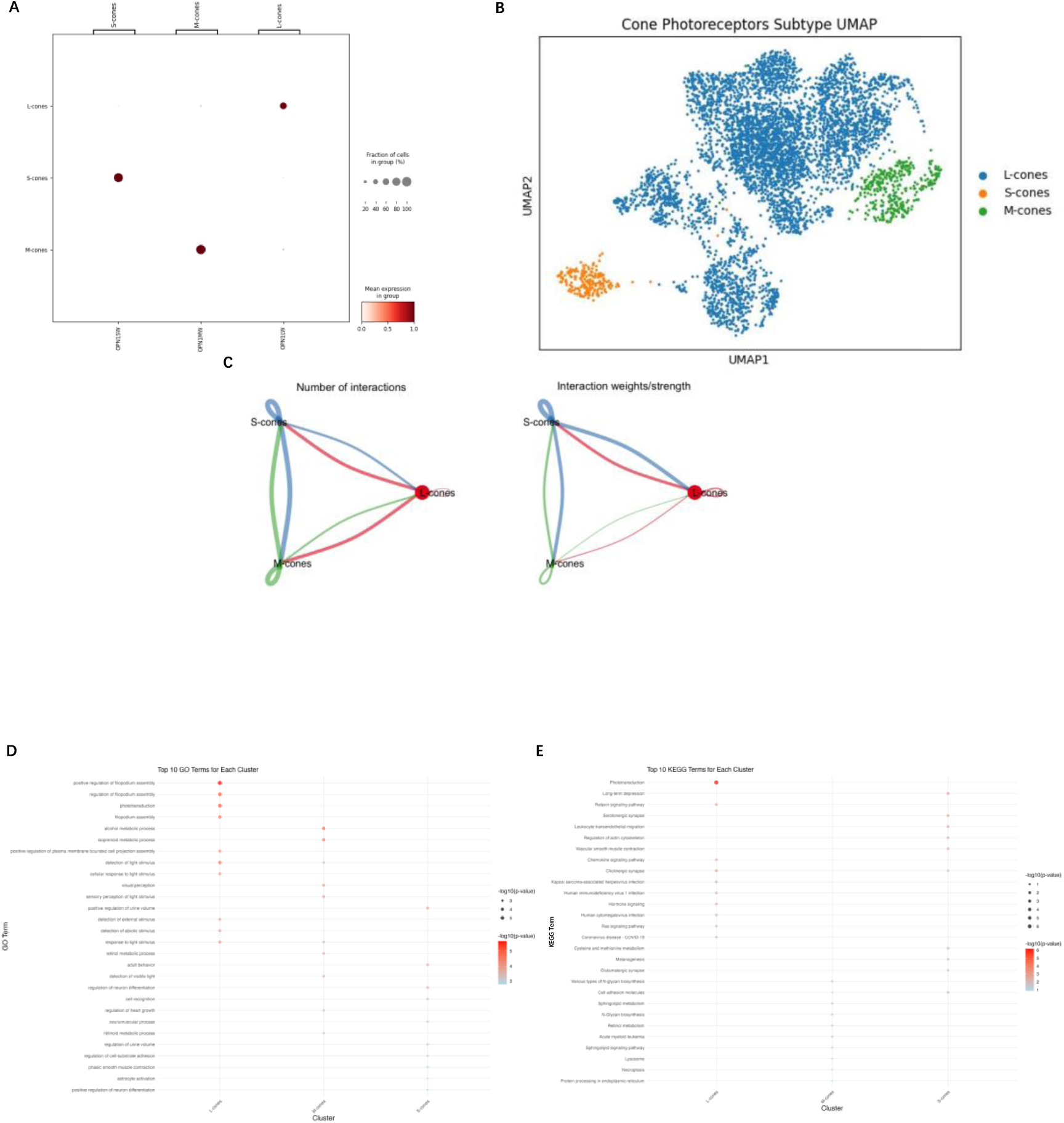
Comprehensive analysis of Chinese human retinal cone photoreceptors type and subtypes. (A) Dot plot demonstrated the identification of each cluster using known markers of subtypes of cone photoreceptors. (B) UMAP plotted that each cell cluster was annotated by cone photoreceptors subtypes. (C) Circle plot visualized the aggregated cell-cell communication network that presented the number of interactions (the left) and the total interaction strength/weights (the right) between any two cell subtypes. (D) Dot plot presented the top 10 enriched GO terms for each cell subtype. The row represented each GO term that these subtypes were involved in. Dots represented that cell subtypes were associated with the GO term, the redder the color, the deeper the correlation. (E) Dot plot presented the top 10 enriched KEGG terms for each cell subtype. The row represented each KEGG term that these subtypes were involved in. Dots represented that cell subtypes were associated with the KEGG term, the redder the color, the deeper the correlation.

The GO and KEGG enrichment analysis of three cone cell subtypes revealed significant biological processes associated with high-acuity vision (detection of light stimulus), phototransduction (phototransduction), cellular morphogenesis (regulation of filopodium assembly), and metabolic regulation (retinol metabolic process), underscoring their specialized roles in visual function (Figure 7D and 7E).

### Rod Photoreceptors

Rod photoreceptors are anther specialized photoreceptor cells located in the human retina. In contrast to cone photoreceptors, rod photoreceptors are predominantly concentrated in the peripheral regions of the human retina [46]. The functions of them are primarily responsible for vision under dark conditions, as they are highly sensitive to light, which enables us to see in low-light environments. Therefore, unlike cone photoreceptors, rod photoreceptors do not contribute to color vision; and they are responsible for monochromatic vision in dark conditions [47]. This is also the reason that we perceive the world in shades of gray in dim light.

Aimed to explore the subtypes of rod photoreceptors, we also performed the cell clustering procedures on them. The UMAP of the cell clustering can be found in the Supplementary Figure 1. Rod photoreceptors have clearly clustered into different cell clusters, but there is no relevant literature describing the cellular subtypes of rod photoreceptors, and we were unable to determine the types of these cell clusters.

### Microglia

Microglia located in the inner and outer plexiform layers are specialized immune cells intrinsic to the central nervous system, playing an essential role in maintaining retinal homeostasis and mediating responses to injury and disease [48]. They are involved in various functions such as immune surveillance, synaptic pruning, and response to injury. Under homeostatic conditions, microglia monitor the retinal environment, remove cellular debris, regulate synaptic remodeling and support neuronal health. In response to retinal diseases, microglia can become activated, releasing inflammatory cytokines, so initial activation may confer neuroprotective benefits but potentially contributing to neuronal damage.

We identified 3 subtypes of microglia that are Egr2 negative M1 microglia, Egr2 positive M1 microglia and M2 microglia [49] (Figure 8B). As for M1 microglia, the known cell markers are *TMEM119* and *EGR2*. The gene *EGR2* is expressed abundantly as the Egr2 positive M1 microglia and vice versa as the Egr2 negative M1 microglia. *CCR5* is the known cell markers for M2 microglia (Figure 8A). Egr2 positive M1 microglia is associated with a robust pro-inflammatory response, which tends to produce higher levels of inflammatory cytokines and activates downstream signaling pathways. Egr2 negative M1 microglia lacking detectable Egr2 expression may be involved in initiating the inflammatory response through mechanisms such as activation of the complement system. M2 microglia display an anti-inflammatory or tissue repair phenotype, which are associated with low levels of pro-inflammatory mediators. The functions of M2 microglia are to maintain human retinal homeostasis and facilitating recovery after injury.

**Figure 8:**
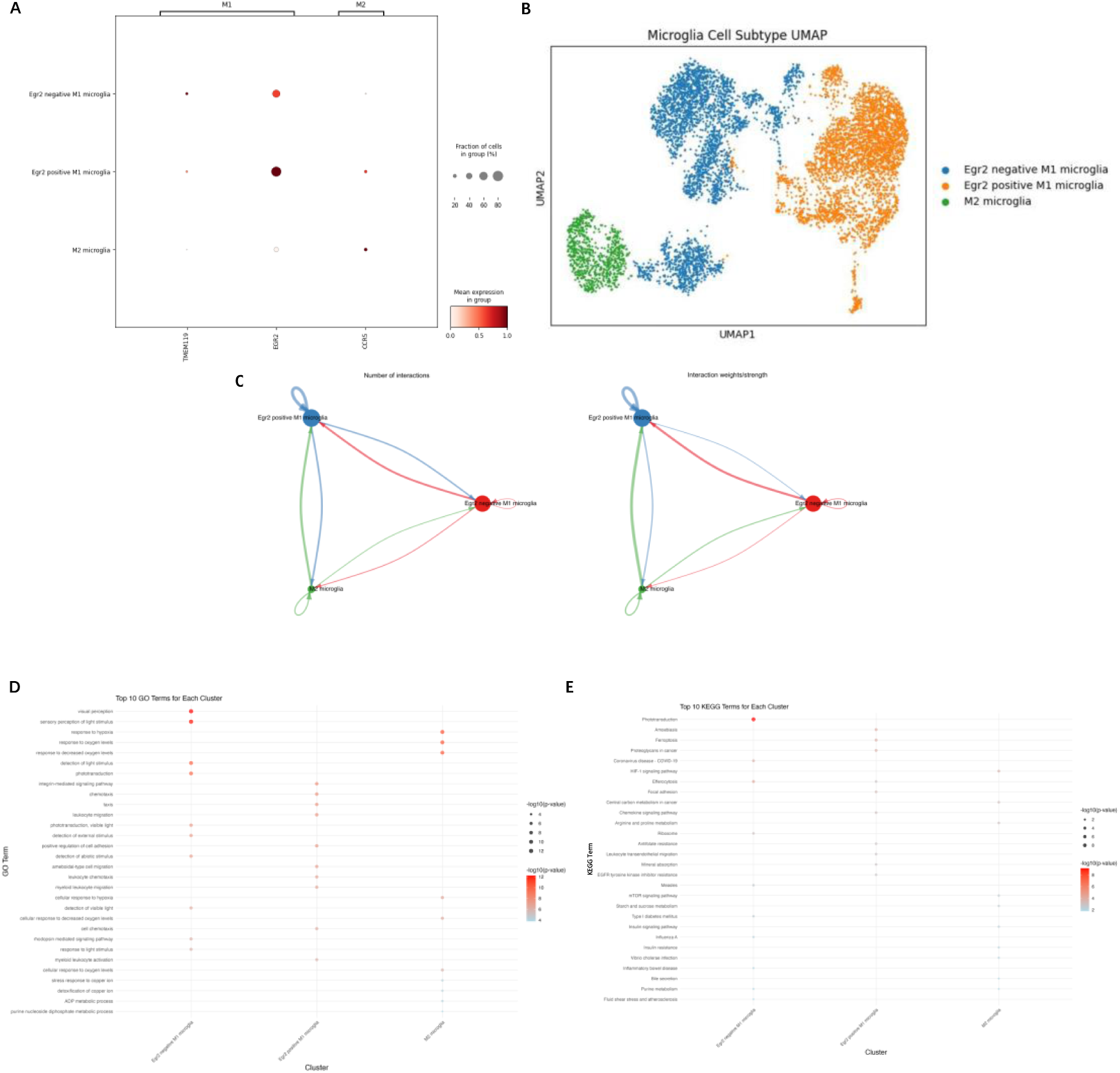
Comprehensive analysis of Chinese human retinal microglia type and subtypes. (A) Dot plot demonstrated the identification of each cluster using known markers of subtypes of microglia. (B) UMAP plotted that each cell cluster was annotated by microglia subtypes. (C) Circle plot visualized the aggregated cell-cell communication network that presented the number of interactions (the left) and the total interaction strength/weights (the right) between any two cell subtypes. (D) Dot plot presented the top 10 enriched GO terms for each cell subtype. The row represented each GO term that these subtypes were involved in. Dots represented that cell subtypes were associated with the GO term, the redder the color, the deeper the correlation. (E) Dot plot presented the top 10 enriched KEGG terms for each cell subtype. The row represented each KEGG term that these subtypes were involved in. Dots represented that cell subtypes were associated with the KEGG term, the redder the color, the deeper the correlation.

The GO and KEGG enrichment analysis of microglia subtypes revealed distinct biological processes linked to phototransduction, visual perception, immune regulation (myeloid leukocyte migration), environmental sensing (detection of external stimulus), and metabolic adaptation (response to hypoxia) (Figure 8D and 8E). Microglia played diverse roles in neuroinflammation and homeostasis in human retina.

### Astrocytes

Astrocytes are a specialized subtype of glial cells that play an important role in the maintenance, development, and functional integrity of the human retina [50]. Astrocytes that are characterized by a stellate structure are predominantly localized in the nerve fiber layer where they closely interact with retinal ganglion cell axons and vascular structures. The functions of them are to contribute to the formation and stabilization of the blood-retinal barrier, which is essential for the regulation of ion homeostasis, neurotransmitter clearance and metabolic support. We performed cell clustering analysis on astrocytes, and the result was shown in the Supplementary Figure 2.

### Müller Glial Cells (MGCs)

Müller glial cells (MGCs) are the principal retinal glial cells in the human retina, playing essential roles in maintaining retinal architecture and function [51–52]. They are involved in several critical processes such as maintaining extracellular ion and water homeostasis, regulating neurotransmitter, supporting synaptic balance, and protecting retinal neurons and photoreceptors. Cell clustering analysis was systematically conducted on MGCs, with cell distributions visualized in Supplementary Figure 3.

### T Cells

T cells predominantly localized at the retinal interface and perivascular regions and engaged in homeostatic immune surveillance or pathological responses in the human retina [53]. It played an important role especially in the common diseases of human retina such as uveitis or age-related macular degeneration. Cell clustering analysis was applied to T cells, which was presented in Supplementary Figure 4.

## Discussion

In this study, we constructed a comprehensive human retinal single-cell transcriptional atlas using retinal samples from Chinese donors, providing a detailed characterization of retinal cell types and subtypes, summarized in the table 2. Through cell clustering and marker gene analysis, we identified transcriptionally distinct subpopulations, revealed potential functional specializations and offered a deeper understanding of cellular functions and interactions.

**Table 2:**
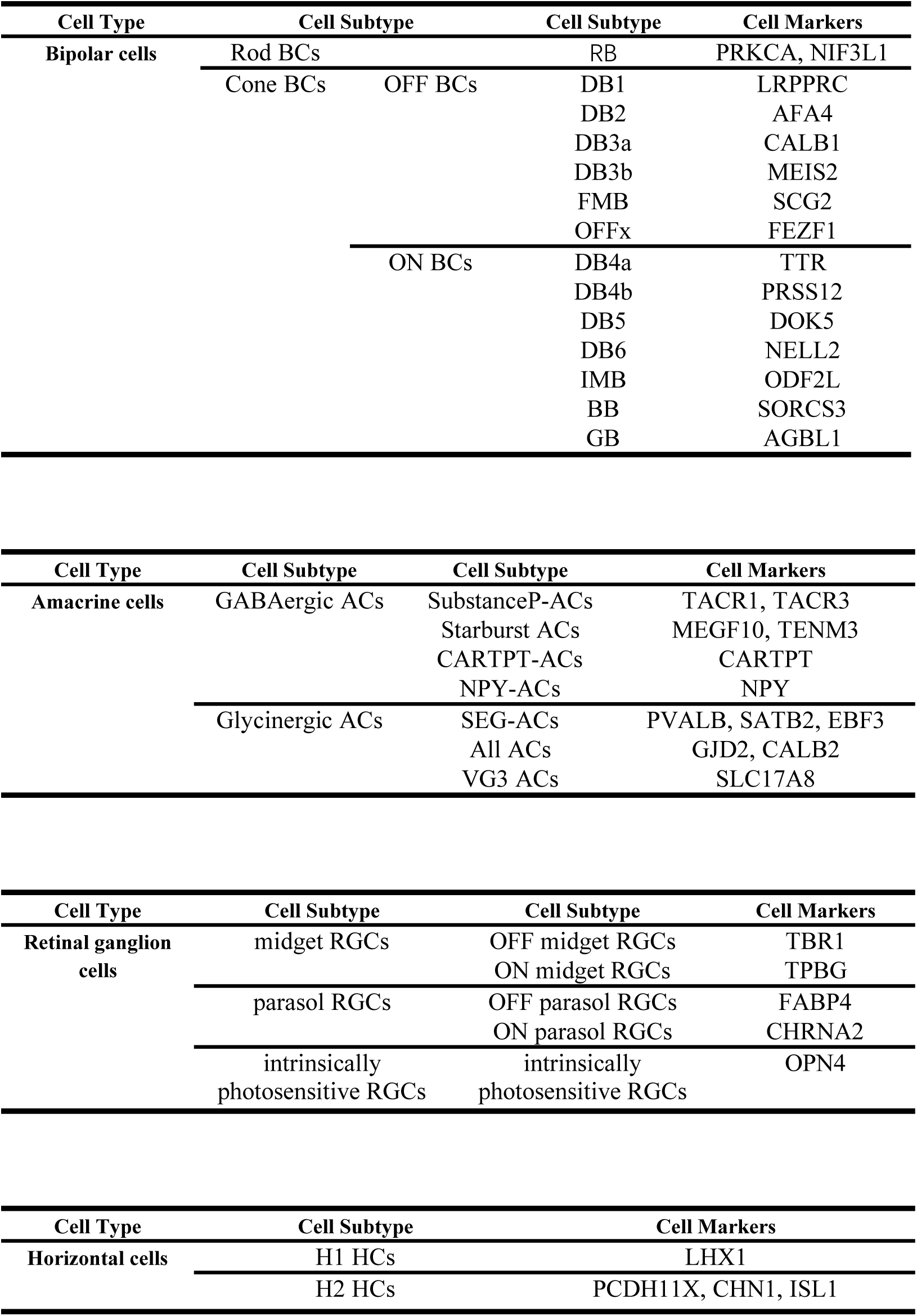
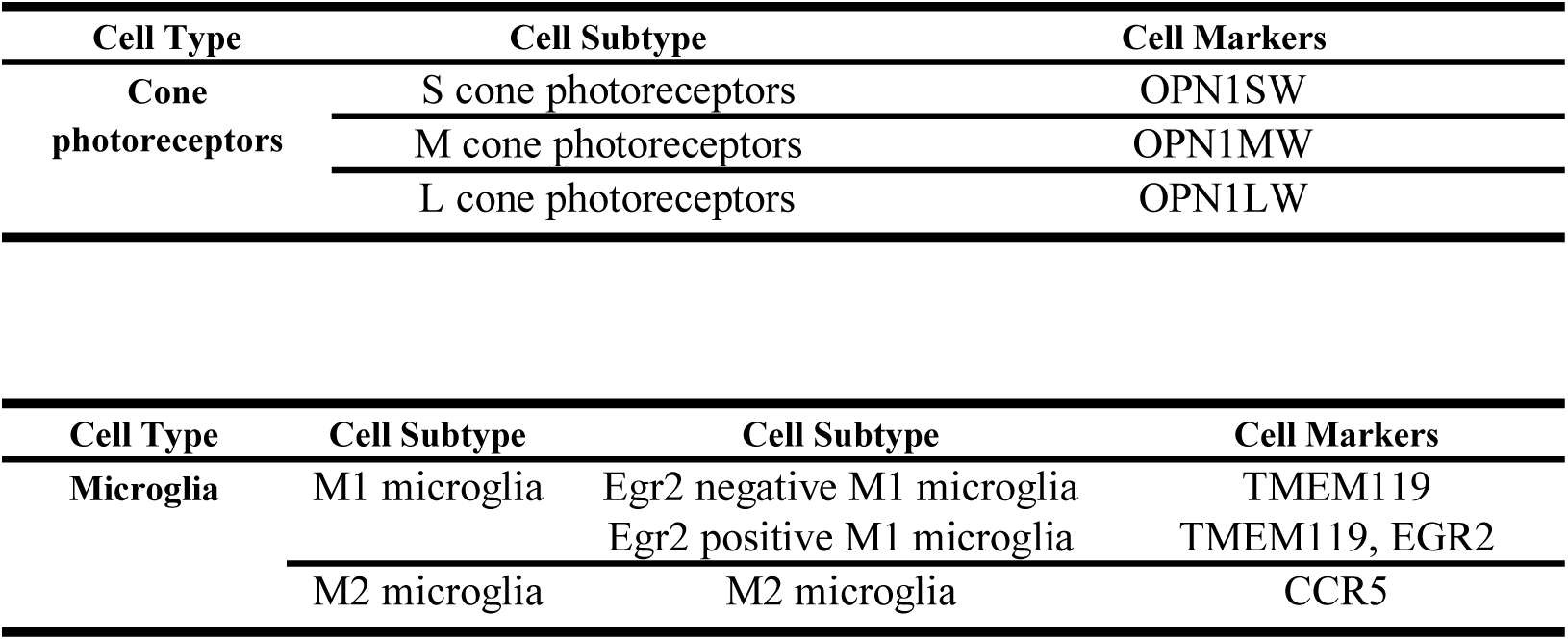
human retinal cell subtype atlas. This table displayed the cell subtypes and cell gene markers in human retina.

| Cell Type | Cell Subtype |  | Cell Subtype | Cell Markers |
| --- | --- | --- | --- | --- |
| Bipolar cells | Rod BCs |  | RB | PRKCA, NIF3L1 |
|  | Cone BCs | OFF BCs | DB1 | LRPPRC |
|  |  |  | DB2 | AFA4 |
|  |  |  | DB3a | CALB1 |
|  |  |  | DB3b | MEIS2 |
|  |  |  | FMB | SCG2 |
|  |  |  | OFFx | FEZF1 |
|  |  | ON BCs | DB4a | TTR |
|  |  |  | DB4b | PRSS12 |
|  |  |  | DB5 | DOK5 |
|  |  |  | DB6 | NELL2 |
|  |  |  | IMB | ODF2L |
|  |  |  | BB | SORCS3 |
|  |  |  | GB | AGBL1 |

| Cell Type | Cell Subtype | Cell Subtype | Cell Markers |
| --- | --- | --- | --- |
| Amacrine cells | GABAergic ACs | SubstanceP-ACs | TACR1, TACR3 |
|  |  | Starburst ACs | MEGF10, TENM3 |
|  |  | CARTPT-ACs | CARTPT |
|  |  | NPY-ACs | NPY |
|  | Glycinergic ACs | SEG-ACs | PVALB, SATB2, EBF3 |
|  |  | All ACs | GJD2, CALB2 |
|  |  | VG3 ACs | SLC17A8 |
| Retinal ganglion cells | midget RGCs | OFF midget RGCs | TBR1 |
|  |  | ON midget RGCs | TPBG |
|  | parasol RGCs | OFF parasol RGCs | FABP4 |
|  |  | ON parasol RGCs | CHRNA2 |
|  | intrinsically photosensitive RGCs | intrinsically photosensitive RGCs | OPN4 |

| Cell Type | Cell Subtype | Cell Markers |
| --- | --- | --- |
| Horizontal cells | H1 HCs | LHX1 |
|  | H2 HCs | PCDH11X, CHN1, ISL1 |
| Cone photoreceptors | S cone photoreceptors | OPN1SW |
|  | M cone photoreceptors | OPN1MW |
|  | L cone photoreceptors | OPN1LW |

| Cell Type | Cell Subtype | Cell Subtype | Cell Markers |
| --- | --- | --- | --- |
| Microglia | M1 microglia | Egr2 negative M1 microglia | TMEM119 |
|  |  | Egr2 positive M1 microglia | TMEM119, EGR2 |
|  | M2 microglia | M2 microglia | CCR5 |

Different gene expression profiles reveal differences in cell types and the dynamics of cell stages. Although previous studies have described some cell types of the human retina, there is still a lack of a cell atlas that describes all major cell types and cell subtypes of the human retina, particularly the lack of a comprehensive analysis of Chinese human retinal samples. In this study, we obtained a relatively complete hierarchical Chinese human retina cell atlas by analyzing retinal cell types and subtypes. This human retinal atlas includes detailed analyses of the cell gene markers of each cell types and subtypes and cellular functions, providing valuable insights into the cellular composition and functional diversity of the Chinese human retina. We collected scRNA-seq from 18 fresh Chinese human retinal samples. Using fresh human retinal samples for analysis was more conducive to accurate cell information extraction.

Our findings align with previous research on retinal cell diversity. From previous literatures, a set of known cellular gene markers was summarized, which can accurately cluster and annotate our human retinal cell data. We identified major human retinal cell types and subtypes in the Chinese donors. GO and KEGG enrichment analysis further systematically infers gene expressions into hierarchical categories of biological processes, molecular functions, cellular pathways and potential metabolic or signaling between cell subtypes. For instance, we found that these cell subtypes of BCs were related to synapse organization and development, regulation of trans-synaptic signaling, and some diseases in the gene level. The CellChat analysis calculated the interactions strength and identifies critical ligand-receptor pairs between cell subtypes. The results demonstrated the key roles of each cell subtype and the detailed biological interactions between these cell subtypes. The discovery of cell subtypes could help elucidate the molecular underpinnings that dictate their specialized roles in transmitting visual information, which is pivotal for advancing our understanding of visual processing and retinal pathologies.

Our study, which constructed a retinal cell atlas using Chinese donors, reveals notable differences when contrasted with retinal cell atlas predominantly derived from Caucasian and Hispanic populations [33]. Specifically for some major cell types, the Chinese retinal atlas identified rod photoreceptors as the predominant cellular constituent, constituting approximately 73.3% of the total retinal cell population. This was followed by BCs (8.9%), ACs (1.5%), and RGCs (0.4%). In contrast, the Caucasian/Hispanic populations exhibited distinct proportions, with rod photoreceptors accounting for 35.5% of total cells, BCs 23.9%, ACs 19.9%, and RGCs 6.8%. The discrepancies may be attributed to two principal factors: firstly, while the comparative study incorporated heterogeneous public datasets that potentially included partial retinal sampling methodologies, our study analyzed the entire human retina. Secondly, the multimodal approach employed in the comparative study (integrating Assay for Transposase-Accessible Chromatin with sequencing (ATAC-seq), scRNA-seq, and single-nucleus RNA sequencing data (snRNA-seq)) contrasts with our exclusive focus on scRNA-seq data. At the level of cellular subtypes, our analysis provided detailed resolution within specific classes—such as identifying 7 ACs subtypes and 14 BCs subtypes—indicating functional specialization within these neuronal circuits. While the comparative study reported equivalent BCs subtype numbers (14 clusters), it described greater ACs diversity (14 clusters) relative to our findings. Furthermore, our analysis extended to non-neuronal cell populations, revealing 3 microglial cells subtypes and 2 HCs subtypes, thereby expanding current understanding of retinal cellular diversity. Quantitatively, the disparity in the number of cells analyzed— 290,401 cells in our study versus about 2 million in the Caucasian/Hispanic work—directly impacts the sensitivity to rare cell types. While our study robustly captures the predominant cellular architecture of the human retina in a Chinese population, the Caucasian/Hispanic dataset’s greater cellular throughput allowed for the identification of less abundant cell populations. These differences underscore the importance of including diverse donor populations in cell atlas projects to reveal population-specific cellular features, which may have implications for understanding ethnic disparities in human retina.

Despite the significant contributions of this study, certain limitations exist. while we have extensively analyzed human retinal cellular types and subtypes, further experimental validation is necessary to confirm the existence of certain cellular subtypes and the functional roles of these cells. For instance, we found the distinct cell clustering results in the rod photoreceptors, but no known literature indicated that rod photoreceptors have cellular subtypes or cellular states. Also, we analyze the functions of cell subtypes based on bioinformatic analysis, not immunohistochemistry. Future research should aim to explore the functional characteristics and interactions of human retinal cells in greater detail. In addition, the study only focused on Chinese populations. Future studies integrating multi-ethnic datasets with standardized protocols will be essential to elucidate gene-environment interactions and establish globally representative retinal reference frameworks for precision ophthalmology.

In summary, the key contribution of this study is to provide a valuable resource for understanding the diversity and functions of retinal cells in Chinese human samples. The constructed atlas enhances our knowledge of retinal cellular composition and offers a basis for future research into retinal diseases and their clinical applications.

## Statements and Declarations

### Data availability

Data will be shared upon publication.

## Authors’ Contributions

SL and YT drafted the paper and performed the analysis. LY, QP, TC contributed to data formatting and correction. YY, JL, YZ, QY, ZL, LC, GM, and RR provided comments on the paper. YS and WM organized the project and provided comments.

## Acknowledgment

This study was mainly funded by the Pioneer and Leading Goose R&D Program of Zhejiang Province 2023 with reference number 2023C04049 and Ningbo International Collaboration Program 2023 with reference number 2023H025.

## Corresponding authors

Correspondence to Yongqing Shao and Weihua Meng.

## Consent to Publish

All authors have consent for publication.

## Ethics approval

This study was approved by the Ethics Committee of the University of Nottingham Ningbo China; The Lihuili Hospital affiliated with Ningbo University; The Affiliated Ningbo Eye Hospital of Wenzhou Medical University.

