## Supplementary figures and images for "Single-cell transcriptomic atlas of human retina from Chinese donors reveals population-specific cellular diversity"

### Supplementary Figure 1

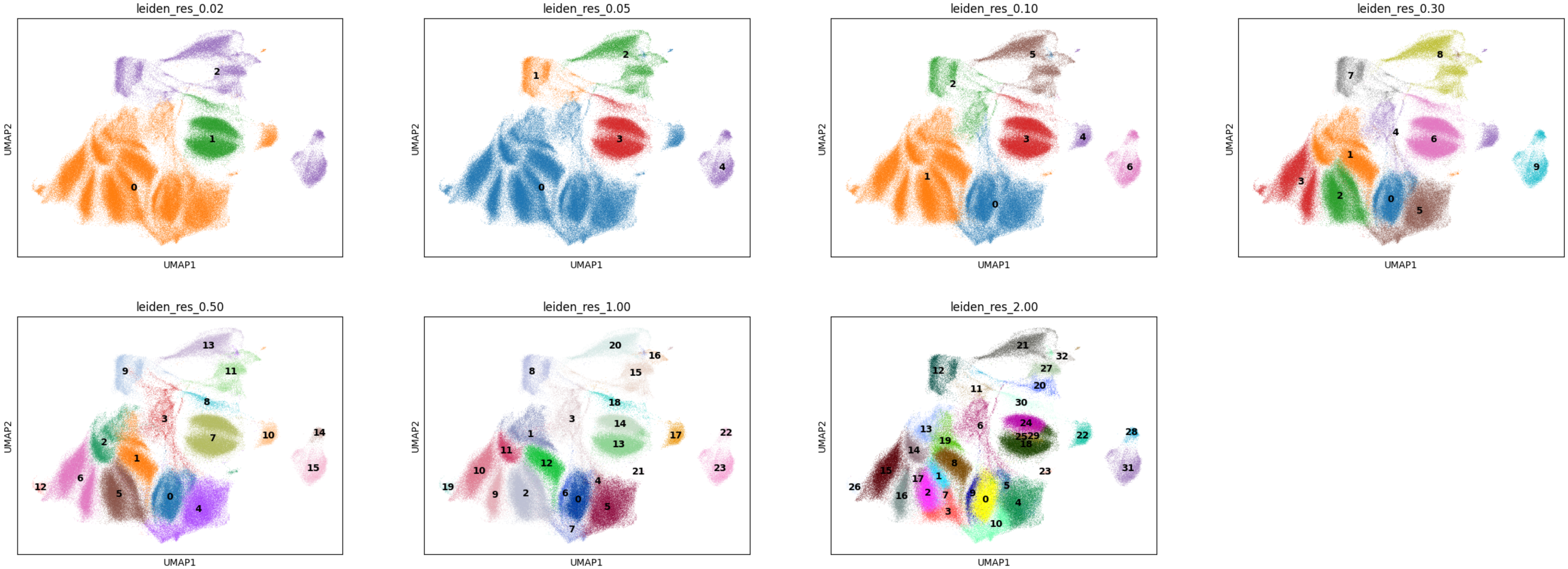

### Supplementary Figure 2

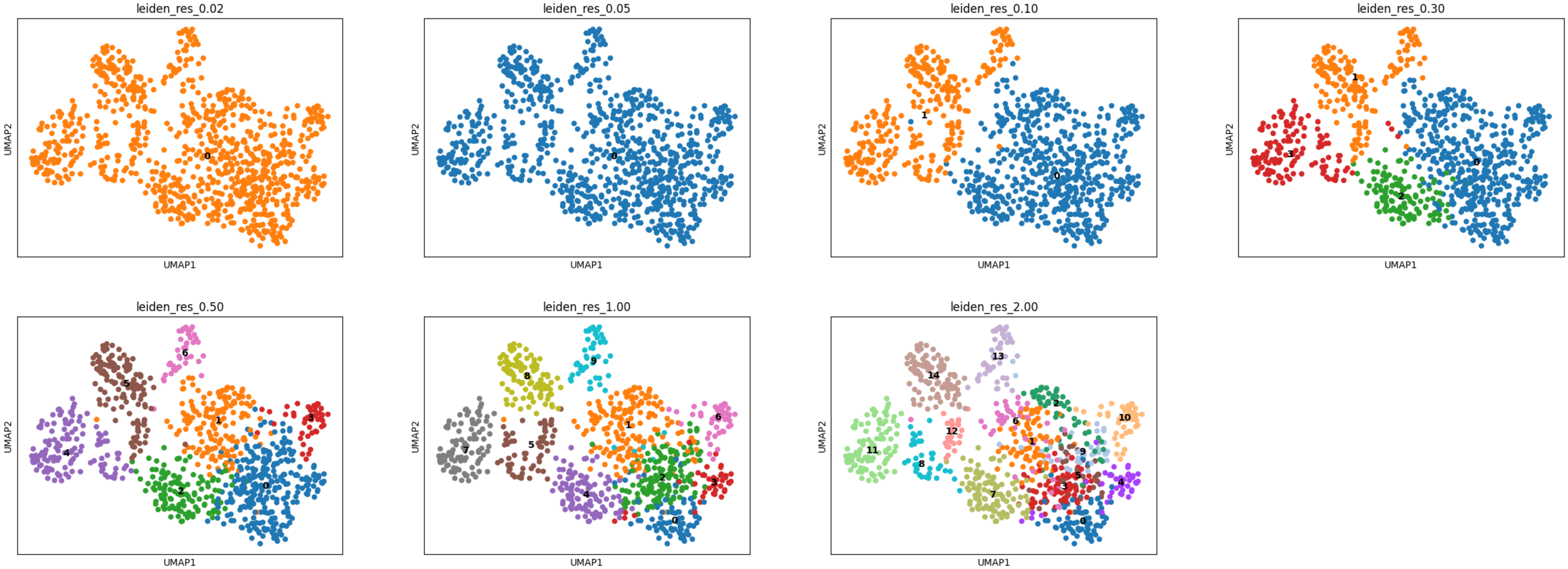

### Supplementary Figure 3

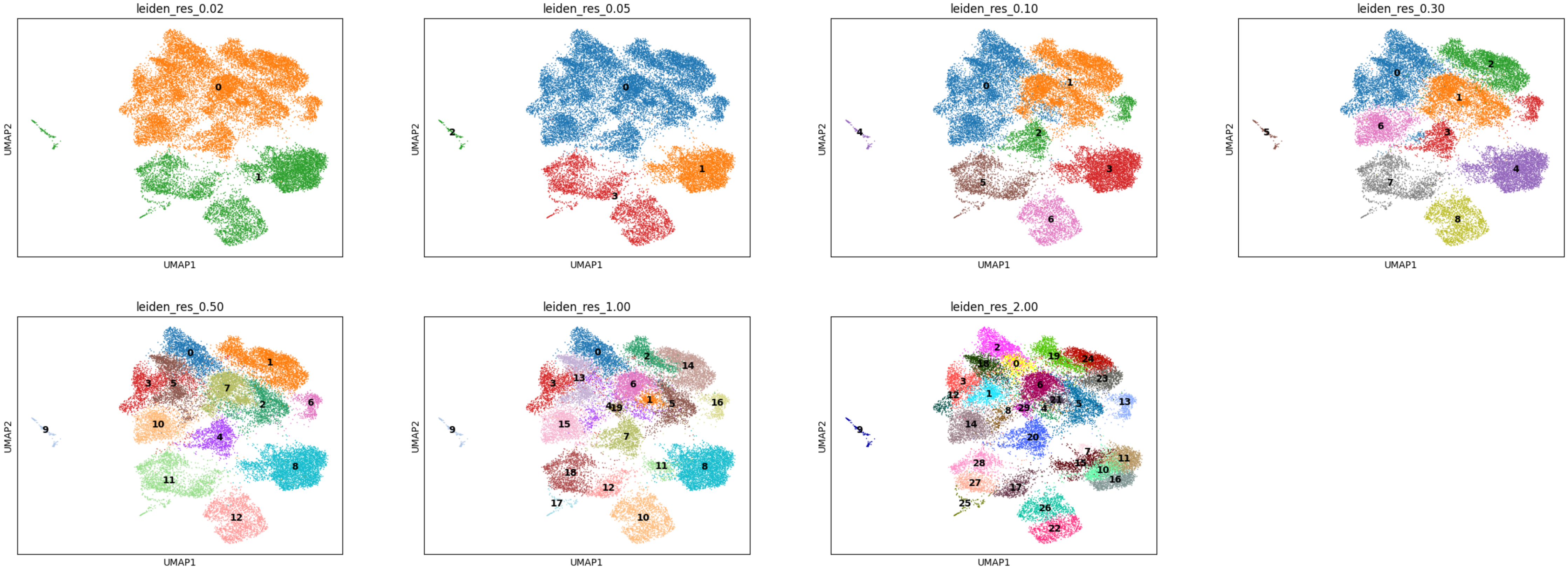

### Supplementary Figure 4

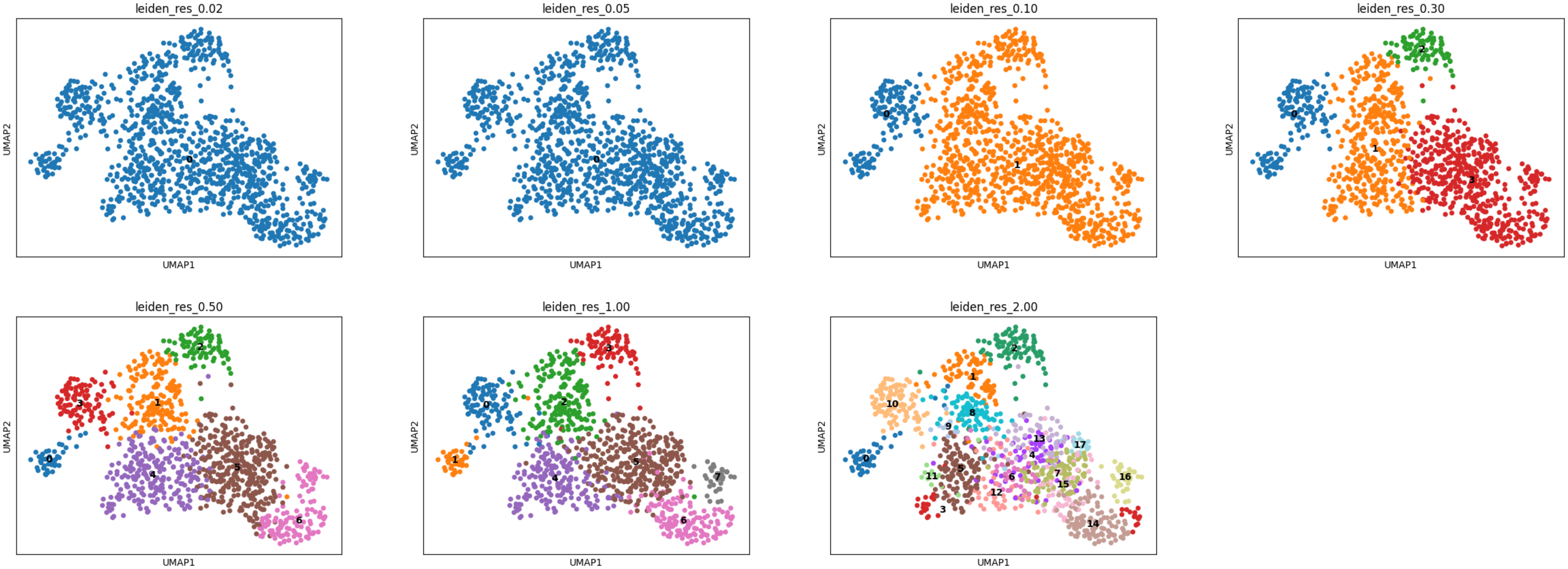
